# Cryo-EM structures of Egl–BicD–RNA complexes reveal how diverse mRNAs are selected for subcellular localization

**DOI:** 10.1101/2025.08.02.668268

**Authors:** Kashish Singh, Sabila Chilaeva, Mark A. McClintock, Andrew P. Carter, Simon L. Bullock

## Abstract

Localization of mRNAs is a widespread mechanism for controlling where proteins operate in cells and underpins many fundamental processes, from embryonic patterning to synaptic plasticity. This spatial control is mediated by RNA-binding proteins (RBPs) that interact with ‘localization signals’ within target mRNAs. However, these signals lack overt sequence or structural patterns, even when targeted by the same protein, raising the question of how specificity is achieved. Here, we investigate this question using the *Drosophila* RBP Egalitarian (Egl), which couples developmentally important mRNAs to microtubule-based transport via Bicaudal D (BicD) and the dynein motor protein. We present cryo-EM structures of Egl– BicD in complex with six different RNAs. Egl uses multiple non-canonical double-stranded RNA-binding domains to cooperatively form a recognition pocket around a localization signal. Despite substantial variation in length and sequence, each signal adopts a bent stem-loop conformation that, together with base-pair identities at two defined sites, drives Egl binding. We further demonstrate that Egl dimers couple RNA binding to transport initiation through coincident detection of two RNA elements within the same transcript. Collectively, we show that diverse localizing mRNAs are recognized through a combination of shape, positional sequence features, and number of structured RNA elements. This work provides a framework for understanding how other RBPs engage highly variable mRNAs.

## Introduction

Localization of mRNAs to specific regions of cells is an evolutionarily conserved strategy for determining where proteins are synthesized and function (*1, 2*). It is essential for numerous biological processes, including embryonic patterning, germline development, establishment of cell polarity, axonal morphogenesis and synaptic plasticity (*3-6*). The main pathways driving mRNA localization are local protection from degradation, hitchhiking on motile organelles, and direct coupling to cytoskeletal motors by adaptor proteins (*2, 3, 7-9*). These mechanisms all rely on the recognition of ‘localization signals’ in mRNAs by RNA-binding proteins (RBPs) (*8, 10*).

Our current understanding of how RBPs identify localizing mRNAs comes largely from cases involving conserved RNA sequences that form linear or structural motifs (*9-12*). However, such clear patterns have proven elusive for most localizing transcripts (*9*). Even in cases where RNA localization signals have been identified and are known to recruit the same RBP, they have considerable divergence in sequence and size (*13-23*). This variability raises the question of how specificity for diverse mRNAs is achieved. The difficulty in identifying localization signals in transcripts has led to suggestions that interactions with RBPs rely on poorly defined structural motifs (*13-21, 24-26*) or combinatorial low-affinity interactions within ribonucleoprotein granules or biomolecular condensates (*27-29*). However, the molecular basis of how such mechanisms can be employed for recognition is not well understood.

An attractive model for investigating how RBPs associate with multiple targets is the Egalitarian (Egl) protein. In the fruit fly *Drosophila melanogaster*, Egl is responsible for patterning and segmentation of the body axes during embryogenesis (*5, 30, 31*), as well as dendritic morphogenesis in larval stages (*32*). It links many different mRNAs to the microtubule motor dynein by binding Bicaudal D (BicD), a long coiled-coil protein with an evolutionarily conserved role in regulating dynein activity (*24, 33*). Association with a target mRNA stabilizes the interaction of two copies of Egl with BicD (*34, 35*). This relieves BicD autoinhibition, leading to activation of dynein movement by the dynactin complex. Many of the RNA localization signals that bind Egl have been defined experimentally (*16-18, 24, 34, 35*). However, besides forming a double-stranded RNA (dsRNA) stem-loop structure, these signals have no obvious similarity in primary sequence or secondary structure (*16-18, 36, 37*). Thus, it is unclear how mRNA recognition is achieved.

## Results

### Architecture of the Egl–BicD complex bound to an RNA localization signal

Egl is a 1004-amino-acid protein in which the first 814 residues are required for specific binding to localization signals (*24*). Alphafold2 (*38*) predicts that the RNA-binding region contains five folded domains connected by flexible linkers (Fig. 1, A and B, and S1A). At the N-terminus are three ‘<u>E</u>gl <u>d</u>omains’ (ED1–3), which adopt a winged helix-turn-helix-like architecture (*39*) that closely resembles the LOTUS domain (named for its presence in Limkain, Oskar, and TDRD5/7 proteins (*40-44*); Fig. S1, B to E). The EDs are followed by a 3′–5′ <u>exo</u>nuclease <u>h</u>omology <u>d</u>omain (ExoHD) (*33*), which is linked to a tetrahelical region that we term the e<u>x</u>onuclease <u>a</u>djacent <u>d</u>omain (XAD). Whereas LOTUS and 3′–5′ exonuclease domains can interact with single-stranded RNA (*42, 45*), there is no evidence they bind dsRNA.

**Fig. 1.**
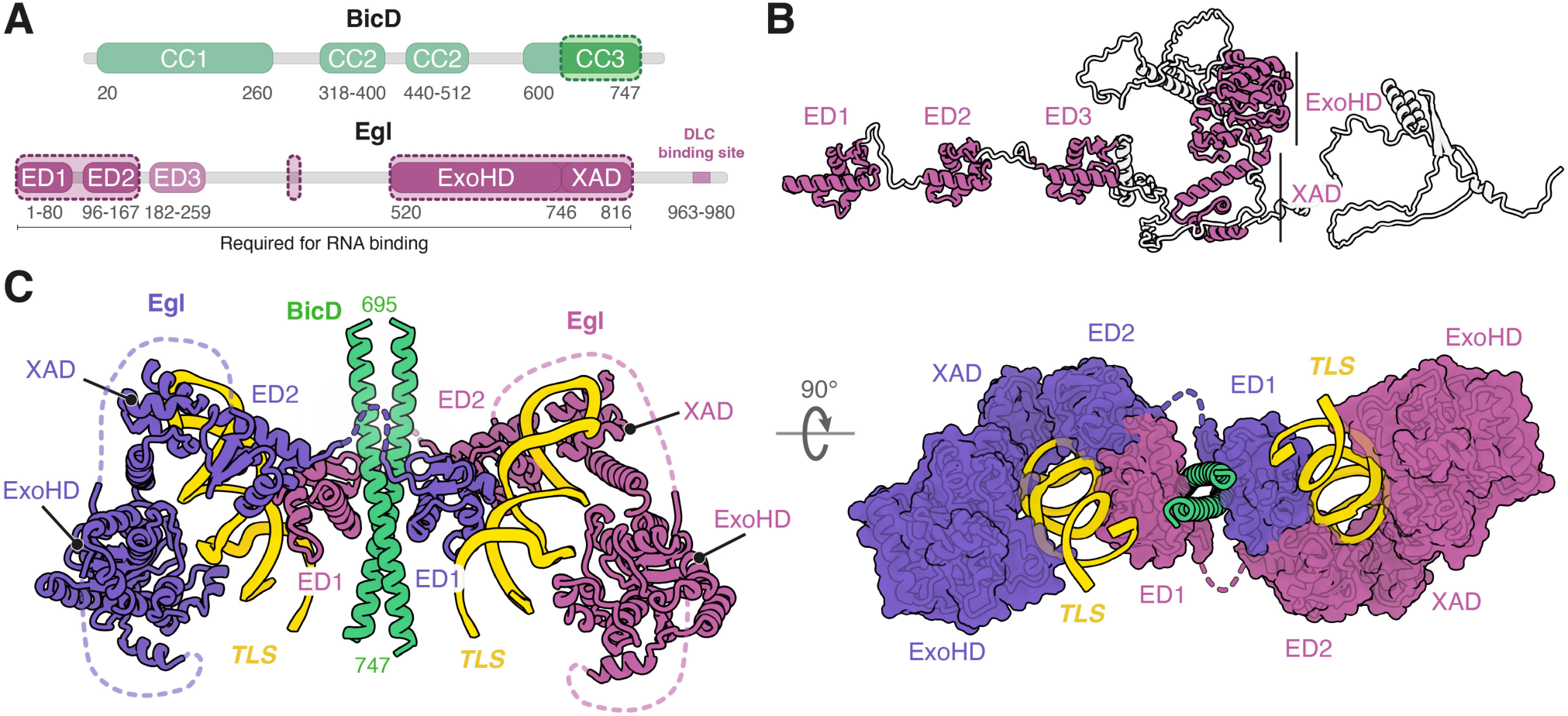
Architecture of the Egl–BicD–*TLS* complex. (**A**) Domain architecture of Egl and BicD. Darker segments enclosed in dashed lines indicate regions visualized in the cryo-EM structure. The Egl C-terminus contains a dynein light chain (DLC)-binding site that promotes Egl function in vivo but is dispensable for dynein activation and RNA binding (*34, 35, 47*). (**B**) Alphafold2 model of Egl with linker regions extended to facilitate visualization of predicted folded domains (colored in magenta; see Fig. S1A for non-extended prediction). (**C**) Cartoon and surface models of the Egl–BicD–*TLS* complex as determined by cryo-EM. Unresolved linkers between domains are represented as dotted lines. The structure was resolved to a nominal resolution of 3.4 Å with local resolution varying from 2.9–4.5 Å (Fig. S2), allowing assignment of RNA and protein sequences to the cryo-EM density (Fig. S3, A and B).

To elucidate how Egl recognizes its dsRNA targets, we first determined a cryo-EM structure of the Egl– BicD complex bound to the 44-nucleotide ‘transport and localization signal’ (*TLS*) of the *Drosophila fs(1)K10* mRNA (*16, 46*) (hereafter *K10*; Fig. S2, S3, A to C, and Table S1). This reveals that two *TLS* stem loops are recruited by two Egl molecules to the C-terminal coiled coil (CC3) of BicD, which is the only region of the latter protein that is resolved in the structure (Fig. 1, A and C). Each copy of Egl binds opposite faces of the BicD coiled coil through ED1 (Fig. 1C, and S3, D and E). ED1 also contacts the *TLS* as part of an RNA-binding pocket that includes an ED2, ExoHD and XAD (Fig. 1C). ED3 is not visible in our structure, indicating it does not stably contact the RNA. Strikingly, within each pocket, the *TLS* is bound by ED1 and ED2 from different Egl polypeptides. This is due to an unstructured loop between ED1 and ED2 that allows the latter domain to contact ED1 of the neighboring Egl molecule but is too short to form intramolecular ED1–ED2 contacts (Fig. 1C). Thus, RNA binding requires two copies of Egl, explaining how localization signals stabilize the dimeric form of Egl that activates transport (*34, 35*).

**Fig. 2.**
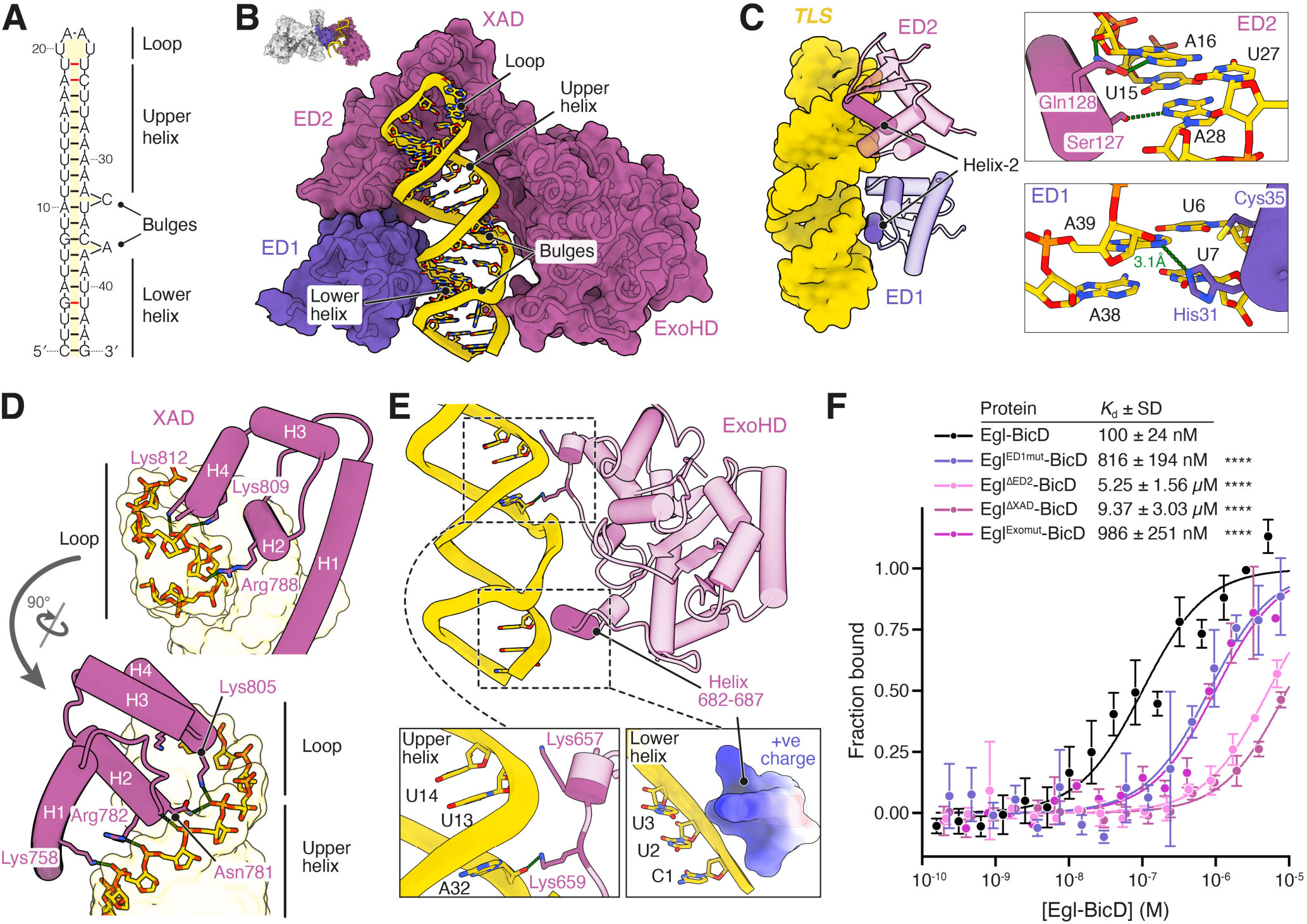
Domains from Egl dimers form composite RNA-binding pockets. (**A**) Empirically determined secondary structure of the *TLS* in complex with Egl–BicD. Non-canonical base pairs are indicated by red lines. (**B**) Overview of the *TLS* in the composite RNA-binding pocket formed from two Egl polypeptides. (**C**) Interaction of ED1 and ED2 with the *TLS* minor groove (left) and side-chain contacts made by these domains with RNA bases (right). His31 of ED1 interacts with the adenine of U6–A39. Ser127 and Gln128 of ED2 interact with the adenines of U15–A28 and A16–U27, respectively. Dotted green lines denote hydrogen bonds. Additional interactions with the sugar-phosphate backbone are shown in Fig. S4, B and C. (**D** and **E**) Interactions of XAD (D) and ExoHD (E) with the *TLS* RNA backbone. Due to limited local resolution of the ExoHD at the lower helix of the *TLS*, a surface charge representation of residues 682–687 is shown in E to indicate likely electrostatic interactions. Additional interactions with the sugar-phosphate backbone are shown in Fig. S4, E and F. (**F**) MST curves for the *TLS* bound to Egl–BicD variants. The Egl^ED1mut^–BicD variant has a Cys35 to Tyr mutation, which is present in the classical *egl^4e^* mutation(*47*). This mutation impairs mRNA association with Egl in vivo(*48*) and is predicted by our structure to clash with nucleotides in the *TLS* minor groove. The Egl^Exomut^–BicD variant has Lys657, Lys659, Arg686 and Arg687 substituted for alanine. Data points show mean ± SD for 3–6 replicates per condition, from which best-fit values for *K*d ± SD were derived. Statistical significance was determined using an extra sum-of-squares F test. ****: p < 0.0001.

**Fig. 3.**
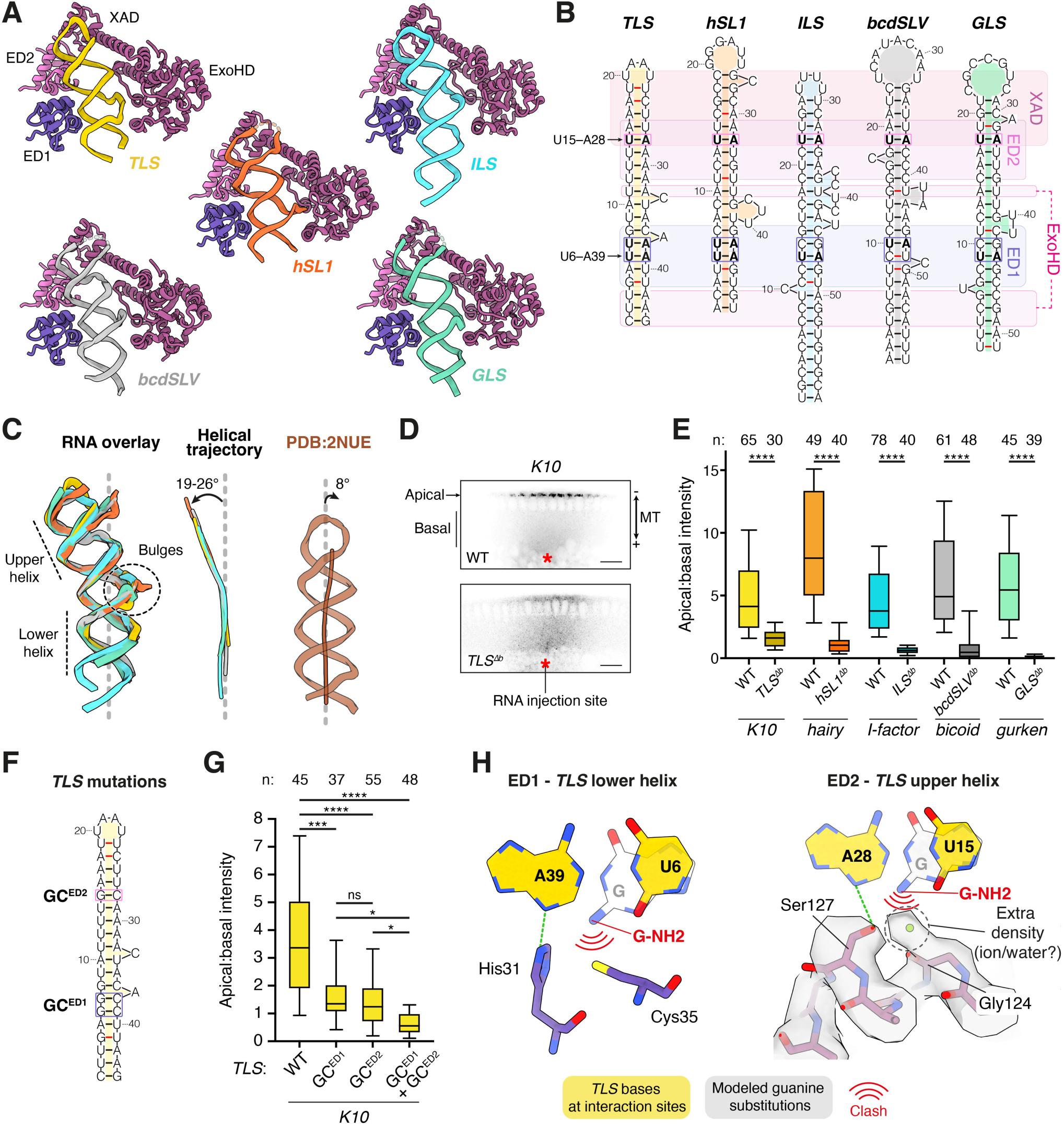
Egl–BicD recognizes shared structural and sequence elements of localization signals. (**A**) Cartoon models of Egl’s composite RNA-binding pocket in complex with the indicated stem loops, as determined by cryo-EM. (**B**) Empirically determined secondary structure of indicated localization signals in complex with Egl– BicD. Stem loops are aligned based on the regions that interact with Egl’s RNA-binding modules (rounded rectangles; purple and magenta differentiate the two Egl polypeptides in the complex). Non-canonical base pairs are indicated by red lines. Base pairs adjacent to ED1 residues His31 and Cys35, and ED2 residue Ser127 are boxed, with the U–A base pairs at these positions shown in bold. (**C**) Superpositions of the stem loops(color-coded as in B) (left) and their helical trajectories (center) compared to a canonical stem loop lacking bulged nucleotides (PDB: 2NUE (*50*); right). (**D**) Confocal Images of *Drosophila* embryos injected with fluorescently-labeled *K10* RNA or a mutant in which the bulges were deleted from the *TLS* (*TLS^Δb^*). Microtubule (MT) polarity and apical and basal regions are indicated. Red asterisk represents RNA injection site. Scale bar, 20 µm. (**E**) Localization efficiency (apical:basal intensity) following injection into *Drosophila* embryos of indicated wild-type fluorescent RNAs and variants lacking bulges in localization signals. (**F**) Predicted secondary structure of the *TLS* with indicated base pair substitutions (boxed). (**G**) Localization efficiency of fluorescently-labeled *K10* RNA and *TLS* variants injected into *Drosophila* embryos. (**H**) Structural basis for guanine exclusion at the interface of ED1 and ED2 with the *TLS* minor groove. Observed positions of *TLS* bases are shown in yellow and modeled positions of guanine substitutions are shown in white. Dotted green lines denote hydrogen bonds. In E and G, boxes show median and interquartile range, and bars show 10^th^–90^th^ percentile range. The number (n) of analyzed embryos from 2–6 independent experiments per transcript is shown. Statistical significance was determined using pairwise two-tailed Mann-Whitney tests (E) or a Brown-Forsythe and Welch ANOVA test with Dunnett’s T3 test for multiple comparisons (G). ns: not significant (p ≥ 0.05); *: p < 0.05; **: p < 0.01; ****: p < 0.0001.

### The structural basis of dsRNA binding by Egl

Each *TLS* comprises a 4-nucleotide loop, two A-form RNA helices (‘upper’ and ‘lower’; Table S2 and S3), and a short intervening helix flanked by 3′-strand bulges (Fig. 2A). ED1 and ED2 bind on one side of the stem loop, contacting the lower and upper helices, respectively (Fig. 2B). An interaction between the EDs positions them to bind adjacent RNA minor groves (Fig. 2C, and S4A). At both sites, the second α-helix (helix-2) of the ED inserts into the minor groove and, together with neighboring residues, makes multiple contacts with the RNA sugar-phosphate backbone (Fig. S4, B and C). In addition, side chains from helix-2 interact with individual bases (Fig. 2C).

The XAD sits in the major groove formed between the *TLS* upper helix and loop, where it makes extensive electrostatic contacts with the RNA backbone (Fig. 2D, and S4, D to G). These interactions would not be possible with a regular A-form RNA helix and require the widened major groove at the tip of the *TLS* stem loop. Furthermore, an interaction between XAD and ED2 sets the distance between the ED2 interaction site and the start of the loop (Fig. S4D). Completing the RNA-binding pocket is the ExoHD, which contacts the opposite side of the *TLS* to ED1–ED2 (Fig. 2B). The putative catalytic residues of the ExoHD (*47*) are ∼15 Å away from the RNA, indicating they do not play a role in *TLS* recognition (Fig. S4H). This is consistent with the dispensability of these residues for RNA binding in vitro (*24*) and Egl function in vivo (*47*). The interaction of the ExoHD with the RNA involves two clusters of positively-charged residues (Fig. 2E, and S4G). Whereas the residues interacting with the upper helix are well-resolved, those contacting the lower helix are not, raising the possibility of dynamic interactions of the ExoHD with this region of the *TLS*.

Microscale thermophoresis (MST) (Fig. S5, A to C) revealed that deleting ED2 or XAD severely reduces binding of Egl–BicD to the *TLS* (Fig. 2F). In contrast, and in line with its absence in our structure, removal of ED3 does not affect *TLS* binding (Fig. S5D). As deleting ED1 or ExoHD was not technically possible (see Materials and Methods), we made point mutations at their RNA-binding interfaces and found these significantly impaired association with the *TLS*, albeit less dramatically than the domain deletions of ED2 or XAD (Fig. 2F).

Collectively, our data show how four non-canonical dsRNA-binding domains of Egl work together to achieve high-affinity RNA binding by engaging distinct features of the *TLS*. Whereas the contacts of these domains with the *TLS* are extensive, the interfaces between ED1, ED2 and the XAD are small and not predicted by Alphafold2 (Fig. S1A, and S4, A and D). These observations support a cooperative mechanism for assembly of the RNA-binding pocket in which the RNA reinforces otherwise weak interdomain interactions.

### Shared structural and sequence features underlie recognition of localization signals

To address how Egl recognizes different localization signals, we additionally determined cryo-EM structures of Egl–BicD bound to the *hSL1, ILS, bcdSLV* or *GLS* RNA stem loops, which are required for dynein-based transport of the *hairy, I-factor, bicoid*, and *gurken* mRNAs, respectively (*17, 18, 24, 37*) (Fig. S6 to S9). These elements were chosen as they are predicted to differ from each other, as well as the *TLS*, in length, loop size, and the placement of bulged nucleotides. Contrary to the earlier hypothesis that Egl has distinct binding modes for different RNA targets (49), our structures show that the protein binds to each localization signal in a similar manner. In all cases, ED1 and ED2 contact adjacent minor grooves with ExoHD and XAD enclosing the rest of the stem loop (Fig. 3, A and B, and S10, A and B). Additionally, the interactions between ED1, ED2 and XAD ensure their binding sites on RNA are equidistant across all stem loops (Fig. 3B).

The structures explain how considerable variability among the different stem loops is tolerated. Variation in loop size is permissible because the XAD interacts only with the portion of the loop adjacent to the upper helix. Bulges on the 5′ strand, present in the *ILS*, *bcdSLV*, and *GLS*, are positioned away from Egl-binding sites and therefore do not interfere with recognition. Finally, deviations in positioning of bulges on the 3′ strand are accommodated by pivoting of the ExoHD–XAD module around the XAD–ED2 interface, allowing ExoHD to shift by up to 15 Å while maintaining contacts with the RNA (Fig. S11A).

Despite the structural variability between localization signals, there are common features that were not apparent from previous secondary structure predictions (Fig. S10B). Like the *TLS*, each RNA contains upper and lower A-form helices of at least six base pairs separated by a short stem segment with bulged nucleotides on the 3′ strand (Fig. 3B, and Table S2 and S3). Additionally, all stem loops exhibit a bend of ∼19–26° between the two helical segments, resulting in a remarkably similar overall RNA backbone conformation (Fig. 3C, and S11, B and C). The RNA bend is important for Egl engagement, as modeling suggests that a straight helix (PDB: 2NUE (*50*)) cannot simultaneously contact ED1 and ED2 without disrupting their interdomain interface, or engage both ends of the rigid ExoHD–XAD module (Fig. S11D). Consistent with these observations, MST measurements showed that the straight dsRNA helix does not detectably associate with Egl–BicD (Fig. S11E). Our structures suggest that the bend in localization signals is instigated by the bulged nucleotides on the 3′ strand (Fig. 3C). This provides a mechanistic explanation for previous data showing the bulged nucleotides in the *TLS* are required for *K10* localization (*51, 52*). Supporting the general importance of a bent helix, we found that deleting bulges from each of the five localization signals strongly impairs Egl–BicD binding in vitro (Fig. S12, A to E), as well as dynein-driven apical localization of mRNAs injected into *Drosophila* embryos (Fig. 3, D and E, and S12, F and G).

As well as a common RNA backbone shape, all five localization signals exhibit similarities in how their minor grooves bind ED1 and ED2. Whereas there is considerable variation in the neighboring RNA sequences, the base pair next to Ser127 of ED2 is a U–A in all structures and those base pairs adjacent to residues His31 and Cys35 in ED1 frequently contain U–As (Fig. 3B). These observations raised the possibility that sequence identity at these two sites contributes to selective RNA recognition. To test this, we introduced U–A to G–C mutations in the *TLS* at both the ED1 (GC^ED1^) and ED2 (GC^ED2^) interaction sites (Fig. 3F). We found that each mutation diminishes Egl–BicD binding in vitro (Fig. S13A) and disrupts *K10* localization in the embryo (Fig. 3G, and S13B), while combining them in the same stem loop has even stronger deleterious effects (Fig. 3G, and S13, A and B).

The pattern of hydrogen bond donors and acceptors in the RNA minor groove is such that only adenine and guanine bases can be discriminated (*53*). Analysis of the ED1 sites across all structures suggests that whereas His31 makes a base-agnostic hydrogen bond to help lock the domain in place, Cys35 is positioned so it would sterically clash with the 2-amino group of guanine (Fig. 3H). This is regardless of whether guanine is located on the 5′ or 3′ strand (Fig. S13C) and is analogous to the RNA recognition mechanism described for dsRBD2 of ADAR2, where a methionine enforces specificity (*53, 54*). Accordingly, guanine is consistently absent at the base pair adjacent to Cys35, which corresponds to U6–A39 in the *TLS* (Fig. 3B). This position is occupied by U–A pairs in all structures except *bcdSLV*, where a non-canonical C–U avoids the clash (Fig. 3B, and S13, D to G). At the ED2 site, while Ser127 plays the equivalent role of His31, there is no direct equivalent to Cys35. However, in all structures resolved to better than 3.5 Å (all except *GLS*-bound Egl–BicD), there is additional electron density at the analogous position (Fig. 3H, and S13, D to G). This likely corresponds to an ion or water molecule that is coordinated by Ser127 and the backbone carbonyl of Gly124 and is positioned such that it would clash with a guanine in the base pair equivalent to U15–A28 of the *TLS*. Thus, the structures show that ED1 and ED2 discriminate against guanine at two sites, which are separated by eight base pairs in each signal (Fig. 3B). Together, our findings reveal that Egl recognizes a localization signal through the combination of a bent dsRNA stem-loop structure and base pair identities at two precisely spaced positions.

### Two RNA stem loops are required for efficient transport

Previous studies showed that single copies of mRNAs associate with Egl–BicD, leading to a model in which one localization signal is sufficient for dynein activation (*34, 35*). However, in our structures, Egl–BicD always binds two stem loops. To reconcile these seemingly contradictory results, we reconstituted motile RNA-protein complexes by assembling dynein, dynactin, BicD and Egl with sub-stoichiometric amounts of Cy3- or Cy5-labeled *TLS* RNAs and visualized movement along microtubules in vitro. If single *TLS* molecules were sufficient for motility, we would detect no colocalization of Cy3 and Cy5 signals in motile complexes (Fig. 4A). In contrast, if two stem loops were needed, half the complexes would contain both fluorophores (Fig. 4A). We found that 44% of motile dynein complexes contain both Cy3 and Cy5, revealing the presence of two *TLS* molecules in the vast majority of cases (Fig. 4, B and C). This indicates that initiation of transport is associated with the assembly of Egl dimers around two separate stem-loop elements. Nonetheless, in agreement with the results of previous studies (*35*), performing the assay with the full-length *K10* revealed only 6% colocalization and therefore a single copy of the RNA in most complexes (Fig. 4, B and C).

**Fig. 4.**
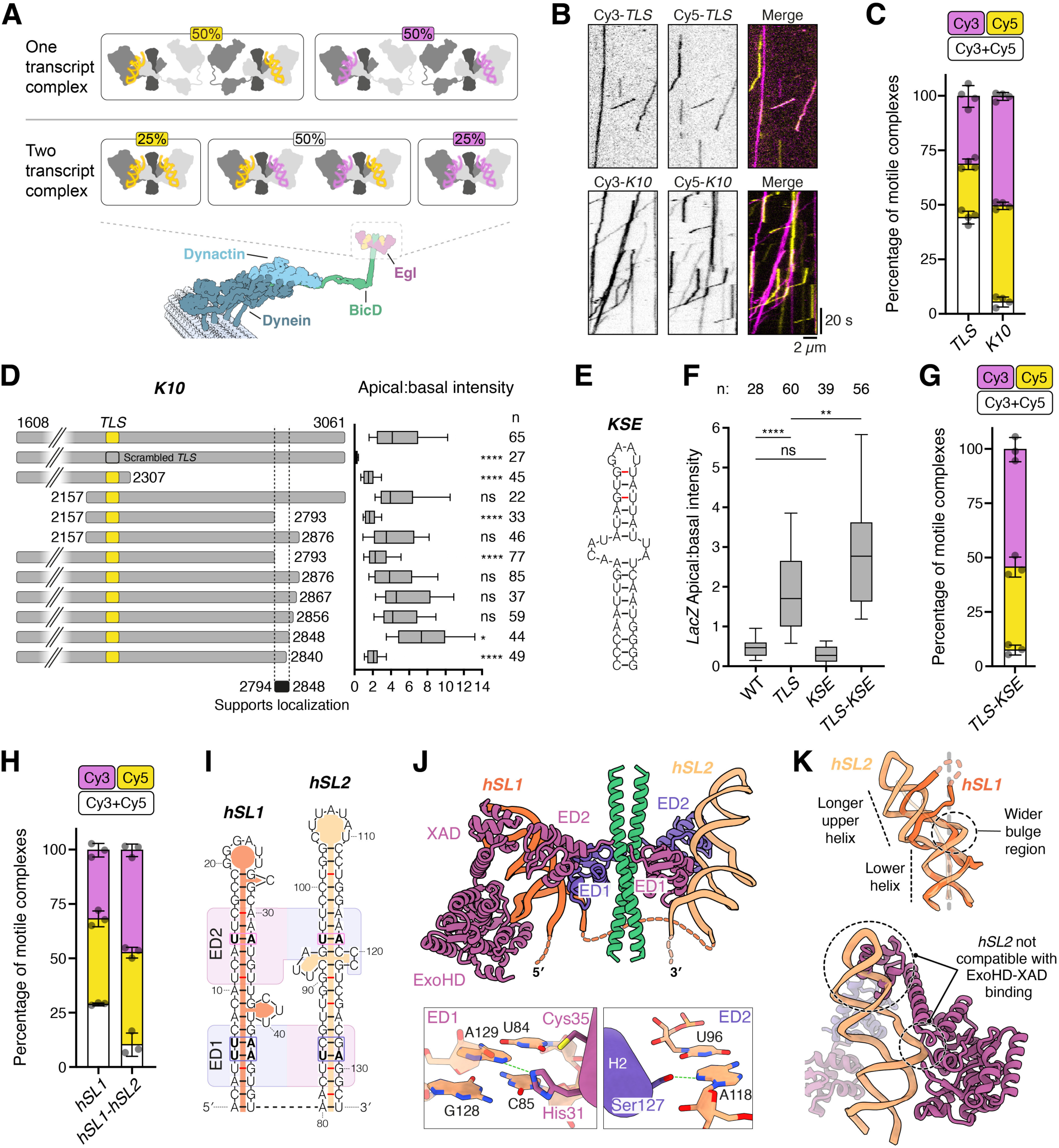
*Cis*-acting support elements promote RNA localization. (**A**) Expected occurrence of fluorescent signals for hypothetical binding of one or two copies of RNA to Egl–BicD in transport complexes reconstituted in the presence of a 50:50 mixture of Cy3- or Cy5-labeled RNA. (**B**) Example kymographs (time-distance plots) from transport assays with a 10-fold molar deficit (relative to Egl) of mixtures of the indicated Cy3- or Cy5-labeled RNAs. (**C**) Observed occurrence of fluorescent signals in motile complexes assembled with the indicated RNAs. (**D**) Truncations of the *K10* 3′ UTR and their localization efficiencies when injected into *Drosophila* embryos. For the intact *K10* 3′ UTR (1608–3061), data are reproduced from Fig. 3E. (**E**) RNAfold-predicted secondary structure of the *KSE* stem loop within the region that supports localization of the *K10* 3′ UTR. (**F**) Localization efficiency of *LacZ* transcripts bearing different combinations of the *TLS* and *KSE* following injection into *Drosophila* embryos. (**G** and **H**) Observed occurrence of fluorescent signals in motile complexes assembled with the indicated RNAs. (**I**) Empirically determined secondary structure of *hSL1* and *hSL2* of the *hairy* localization element in complex with Egl–BicD. Stem loops are aligned based on the regions that interact with ED1 and ED2 (rounded rectangles; purple and magenta differentiate the two Egl polypeptides in the complex). Non-canonical base pairs are indicated by red lines. Base pairs adjacent to ED1 residues His31 and Cys35, and ED2 residue Ser127 are boxed, with the U–A base pairs at these positions shown in bold. (**J**) Cryo-EM structure of Egl–BicD bound to *hSL1* and *hSL2* (top). The unresolved linker between *hSL1* and *hSL2* is depicted as a dotted line. His31 of ED1 and Ser127 of ED2 contact guanine-deficient U84–A129 and U96–A118 base pairs, respectively (bottom). Dotted green lines denote hydrogen bonds. (**K**) Top: Superposition of *hSL1* and *hSL2* stem-loop structures. Bottom: ExoHD–XAD modeled at the *hSL2* binding site showing clashes with the *hSL2* stem loop. The ExoHD–XAD from the *hSL1* binding pocket was overlayed onto the *hSL2* binding site by aligning the two binding pockets using their respective ED2’s. In C, G and H, black dots represent values from 3 or 4 independent experiments and error bars show SD; 235–1,032 motile complexes analyzed per transcript. In D and F, boxes show median and interquartile range, and bars show 10^th^–90^th^ percentile range. The number (n) of analyzed embryos from 2–6 independent experiments per transcript is shown. Statistical significance was determined using a Brown-Forsythe and Welch ANOVA test with Dunnett’s T3 test for multiple comparisons. ns: not significant (p ≥ 0.05); *: p < 0.05; **: p < 0.01; ****: p < 0.0001.

These results could be explained by the full-length *K10* mRNA containing a second, previously unidentified RNA element that binds Egl–BicD along with the *TLS*. To test if *K10* contains a second element that is important for localization, we injected a series of *TLS*-bearing truncations of the RNA into *Drosophila* embryos (Fig. 4D, and S14A). These experiments identified a 54-nucleotide region of the 3′ UTR that is required for efficient apical localization and contains a 40-nucleotide stem loop (Fig. 4, D and E, and S14A). We refer to this structure as the *<u>K</u>10* localization <u>s</u>upport <u>e</u>lement (*KSE*). While the *KSE* cannot promote apical localization of a heterologous *lacZ* RNA on its own, it significantly enhances *TLS*-mediated transport of this transcript (Fig. 4F, and S14B). MST revealed that the *KSE* binds Egl–BicD, albeit with lower affinity than the *TLS* (K_d_ = 507 nM vs. 80 nM; Fig. S14C). We also found using the dual-color motility assay that combining the *TLS* and the *KSE* in a minimal 145-nucleotide RNA results in activation of dynein by a single RNA (Fig. 4G). Therefore, despite the lower independent affinity of the *KSE,* Egl–BicD shows a preference for binding both elements in a single RNA over two *TLS* elements in separate RNAs. This suggests that avidity conferred by the presence of the *TLS* and the *KSE* on the same transcript enhances Egl–BicD association.

The requirement for two RNA elements to activate dynein could explain the reported multipartite nature of certain other localization elements (*18, 37, 55*). To test this possibility, we turned our attention to the 126-nucleotide *hairy* localization element (*18*), which consists of *hSL1* and a second stem loop, *hSL2*. The latter element is not active in isolation but, through an unknown mechanism, stimulates apical localization driven by *hSL1* (*18*). We found that Egl–BicD associates with *hSL2* but with a lower affinity than observed for *hSL1* (K_d_ = 603 nM vs. 7 nM; Fig. S14D). Moreover, dual-color motility assays revealed that while two copies of an isolated *hSL1* are typically needed to activate dynein, an RNA containing both *hSL1* and *hSL2* is much more likely to be transported as a single copy (Fig. 4H). These observations indicate that *hSL2* serves an equivalent function to the *KSE* during activation of the motor, i.e. cooperating with a primary signal to occupy two RNA-binding sites within Egl–BicD. Therefore, our data reveal that the presence of two Egl-binding stem loops on the same RNA is another criterion by which mRNAs are selected for localization.

### The structural basis of dual stem-loop recognition

To understand how the combination of a primary localization signal and support element are recognized, we determined a cryo-EM structure of Egl–BicD bound to the *hairy* localization element (Fig. 4, I and J, and S15). This revealed that *hSL1* is bound by ED1, ED2, ExoHD and XAD (Fig. 4J), as seen in the structure of this stem loop in isolation (Fig. 3A). In contrast, on the opposite side of BicD, *hSL2* only engages ED1 and ED2 (Fig. 4J). The interaction of *hSL2* with these domains is through a bent helical backbone and guanine-deficient base pairs (Fig. 4, I to K, and S16, A and B), as observed for all five primary localization signals (Fig. 3). However, *hSL2* contains additional unpaired nucleotides on the 5’ strand that expand the bulge region, resulting in a larger bending angle and a shift in the position of the upper helix relative to that of *hSL1* (Fig. 4K, and S16, C and D). These structural deviations are accommodated by pivoting of ED2 around its interaction site with ED1, which strains the ED1–ED2 interface but still maintains interactions of the domains with the RNA (Fig. S16E). Additionally, *hSL2* has a longer upper helix than *hSL1*, which, coupled to the wider bulge region, sterically prevents docking of ExoHD– XAD (Fig. 4K). The absence of ExoHD–XAD engagement, as well as the suboptimal arrangement of ED1–ED2, likely explains the relatively weak binding of the isolated *hSL2* to the Egl–BicD complex (Fig. S14D). Overall, this structure reveals how a single transcript engages Egl–BicD through simultaneous association of high- and low-affinity RNA elements.

## Discussion

Our current understanding of RNA recognition largely derives from studies of proteins that target specific sequence motifs or conserved secondary structures (*56, 57*). However, the mechanisms by which RBPs identify RNA elements lacking such well-defined patterns remain poorly understood. Here, structures of Egl bound to different localization signals reveal multiple layers of recognition that govern specific engagement in the absence of obvious conservation of sequence or secondary structure. Egl recognizes a bent dsRNA stem-loop structure that can be formed by diverse sequences containing bulges of varying number and size. In conjunction, Egl discriminates against guanine bases at two distant base-pair positions in the minor groove. Specificity is further increased by the coincident recognition of two stem-loop elements within the same transcript by Egl dimers. Together, tertiary RNA structure recognition, minor-groove base discrimination, and dual-element detection provide a versatile yet robust strategy for selective recognition of diverse mRNA cargoes.

A previous NMR structure of the isolated *TLS* suggested that an A′-form dsRNA conformation contributes to mRNA localization (*52*). Although we did not observe this feature in any of our structures, we noticed that the isolated *TLS* structure is already bent and requires only minor adjustments to adopt the Egl-bound state (root-mean-square deviation (RMSD) = 3.8 Å; Fig. S17, A and B). This contrasts with another structurally characterized RBP–localization signal complex (*58*), where the yeast *ASH1* mRNA element displays a large conformation switch between free and She2p–She3p-bound states (RMSD = 16.3 Å; Fig. S17C). These comparisons suggest that localization signals need not be highly dynamic as previously proposed (*58*) but can instead pre-exist in a stable, recognition-competent conformation.

Another key insight from our study is that the binding of Egl–BicD to two stem loops is a prerequisite for activation of RNA transport. We show that for *K10* and *hairy* this requirement is fulfilled by the combination of a primary localization signal and a support element within the same transcript. This strategy enhances specificity for cargo while also incorporating a lower-affinity interaction that may facilitate efficient recycling of transport complexes after delivery. It can also explain why other Egl target mRNAs – such as *wingless* (*55*), *fushi tarazu* (*37*), and *bicoid* (*37*) – require regions beyond a single stem loop for proper localization. We therefore propose that occupancy of Egl–BicD by two elements within the same RNA is a widespread mechanism for cargo discrimination. However, it is conceivable that some RNAs lack additional support elements and compensate by forming oligomeric structures (*59-61*) that would expose multiple primary localization signals for Egl–BicD engagement, thereby allowing selective transport of higher-order RNA assemblies.

Recognition of tens to hundreds of distinct mRNAs have been demonstrated for other RBPs (*62*). These include factors that orchestrate mRNA localization, such as Staufen2 (*20, 63*), FMRP (*21, 64*), APC (*25, 65*), She2p–She3p (*15, 58*), and the recently characterized FERRY complex (*26, 66*). Similar to Egl, many of these RBPs contain multiple RNA-binding domains and/or form dimeric assemblies and therefore have the potential to recognize diverse RNA elements using multiple selection criteria. However, because identifying RNA recognition motifs has remained challenging, several of these proteins have been suggested to bind their targets via non-specific binding strategies such as assembling into RNA granules or biomolecular condensates (*9, 67-69*). While these models remain plausible, our visualization of Egl– BicD bound to many targets reveals that conserved recognition features can exist even among RNAs that differ substantially in sequence and secondary structure. This raises the possibility that mRNAs targeted by other RBPs also contain currently elusive patterns that support their selective recognition.

## Methods

### Recombinant protein expression

The *Drosophila* Egl–BicD and human dynein complexes were expressed recombinantly from baculovirus in *Sf*9 insect cells using a polycistronic MultiBac system (*70*), as described previously (*34*). Briefly, a codon-optimized coding sequence for Egl (isoform B; NM_166623) with a C-terminal TEV-ZZ affinity tag was synthesized (Epoch Life Sciences) and cloned into the pACEBac1 acceptor plasmid and Cre-recombined with a pIDC donor plasmid containing a codon-optimized coding sequence for BicD (NM_165220; Epoch Life Sciences). The recombined plasmid was incorporated into the baculovirus genome by transformation of DH10EMBacY cells. This baculovirus genome was purified and used with FuGENE HD (Promega) to transfect adherent *Sf*9 cells. After ∼96 hours at 27 °C, transfection/infection of the majority of *Sf*9 cells was confirmed by detection of YFP. The baculovirus was then amplified by using the supernatant from the transfection to infect a larger 50-mL suspension culture of *Sf9* cells for ∼96 hours at 27 °C. The amplified virus was harvested by spinning the 50-mL culture and decanting the virus-containing supernatant, which was subsequently used to infect 500-mL suspension cultures for final protein expression. Cells from these cultures were pelleted, flash frozen in liquid nitrogen, and stored at -80 °C until processed for protein purification. All mutants of Egl were derived from the construct in pACEBac1 describe above by site-directed mutagenesis (Egl^ED1mut^; Cys35Tyr) or by Gibson Assembly (New England Biolabs) of PCR amplicons that excluded the targeted coding sequence (Egl^ΔED2^; deletion of residues 94–167, Egl^ΔED3^; deletion of residues 187–259, Egl^ΔXAD^; deletion of residues 744–816) with the exception of Egl^Exomut^ (Lys657Ala, Lys659Ala, Arg686Ala, Arg687Ala), which was commercially synthesized (Epoch Life Sciences) and cloned into pACEBac1 for use as above. The relative instability of an Egl–BicD complex lacking the ExoHD resulted in insufficient protein concentrations for use. Deletion of ED1 was not pursued as it is expected to compromise Egl stability by eliminating its association with BicD (*34*). The human dynein complex was expressed in an analogous way to Egl–BicD, with sequences encoding Dynein-1 heavy chain (NM_001376.4) bearing an N-terminal ZZ-TEV-SNAPf tag cloned into pACEBac1 and the remaining subunits (Dynein intermediate chain 2 (DIC2; AF134477), Dynein light intermediate chain 2 (DLIC2; NM_006141.2), and the light chains (Tctex; NM_006519.2, LC8; NM_003746.2, and Robl; NM_014183.3)) cloned into pIDC (*71*).

### Protein purification

The Egl–BicD complex (including its variants) and the dynein complex were produced as described previously (*34*), with all purification steps occurring at 4 °C. *Sf*9 insect cells were routinely confirmed as Mycoplasma-free using the MycoALERT kit (Lonza). Cells expressing the recombinant complexes were suspended in lysis buffer (50 mM HEPES pH 7.3, 500 mM NaCl (100 mM NaCl for dynein), 10% glycerol, 1 mM DTT, 0.1 mM MgATP, 2 mM PMSF, and 1X cOmplete EDTA-free protease inhibitor cocktail (Roche)) and lysed by passage in a Dounce homogenizer with a tight pestle. The lysate was clarified by ultracentrifugation in a Type 70 Ti rotor (Beckman-Coulter) at ∼500,000 x g, applied to pre-washed (in lysis buffer) IgG-Sepharose FF resin (Cytiva) in a gravity-flow Econo column (Bio-Rad) and incubated with gentle rolling for 3 hours. Flow-through was collected by gravity and the protein-bound resin was washed twice with five column volumes of lysis buffer and twice with five column volumes of TEV buffer (50 mM Tris pH 7.4, 150 mM KOAc, 2 mM MgOAc, 1 mM EGTA-KOH pH 7.5, 10% glycerol). The resin was then transferred to a 15-mL conical tube in 15 mL final volume of TEV buffer and incubated overnight at 4 °C with TEV protease to cleave the protein complexes from the beads and ZZ affinity tag. The liberated protein complexes were collected by gravity flow from a fresh Econo column and concentrated to a volume of ∼500 µl (Egl–BicD) or ∼300 µl (dynein) using Amicon Ultra-4 100 kDa MWCO concentrator units (Merck) before application to Superose 6 Increase 10/300 (Egl–BicD; Cytiva) or TSKgel G4000SWxl with guard (dynein; TOSOH Bioscience) gel filtration columns run in GF150 buffer (25 mM HEPES pH 7.3, 150 mM KCl, 1 mM MgCl_2_, 0.1 mM MgATP, 5 mM DTT, 10% glycerol) on an AKTA Purifier FPLC instrument (Cytiva). Fractions containing the protein complexes were pooled and concentrated using the same concentrator units described above to final concentrations of ∼1.5 mg/mL (Egl–BicD and dynein for single-molecule motility assays) or 4–5 mg/mL (Egl– BicD for MST assays), as determined by the Bradford assay (Pierce).

The native dynactin complex was purified at 4 °C from pig brain extracts as described previously (*71*). Three pig brains were homogenized in lysis buffer (35 mM PIPES-KOH pH 7.2, 5 mM MgSO_4_, 1 mM EGTA-KOH pH 7.5, 0.5 mM EDTA pH 7.4, 1 mM DTT, 2 mM PMSF, 1X cOmplete EDTA-free protease inhibitor cocktail (Roche)) using short bursts in a blender. The lysate was clarified first at low speed at 38,400 x g in a JLA 16.250 rotor (Beckman-Coulter) and then at higher speed by ultracentrifugation at ∼160,000 x g. The clarified lysate was filtered through a 0.45-µm syringe-tip filter (Fisher) and loaded onto an SP-Sepharose (Cytiva) cation exchange column (∼250 mL bed volume) equilibrated in SP-Buffer A (35 mM PIPES-KOH pH 7.2, 5 mM MgSO_4_, 1 mM EGTA-KOH pH 7.5, 0.5 mM EDTA pH 7.4, 1 mM DTT, 0.1 mM MgATP) and run on AKTA Pure FPLC instrumentation. The lysate was fractionated by washing the column with six column volumes of Buffer A and eluting with a two-phase linear salt gradient with SP-Buffer B (SP-Buffer A with 1 M KCl) in which the first phase increased the KCl concentration to 250 mM over three column volumes and the second phase further increased the KCl concentration to 1 M over an additional one column volume. Dynactin-containing fractions were pooled and diluted two-fold with Q-Buffer A (35 mM PIPES-KOH pH 7.2, 5 mM MgSO_4_, 1 mM EGTA-KOH pH 7.5, 0.5 mM EDTA pH 7.4, 1 mM DTT) and filtered through a 0.22-µm syringe-tip filter (Fisher) before loading onto a MonoQ 16/10 column (Cytiva) equilibrated in Q-Buffer A, and running on AKTA Pure FPLC instrumentation. After loading, the MonoQ 16/10 column was washed with five column volumes of Q-Buffer A and the bound protein eluted in a three-phase linear salt gradient with Q-Buffer B (Q-Buffer A with 1 M KCl) in which the first phase increased the KCl concentration to 150 mM over one column volume, the second phase increased the KCl concentration to 350 mM over 10 column volumes, and the third phase increased the KCl concentration to 1 M over one column volume. Fractions containing dynactin were pooled and concentrated in an Amicon Ultra-4 100 kDa MWCO concentrator unit (Merck) to a final volume of 300–500 µl before application to a TSKgel G4000SWxl with guard (TOSOH Bioscience) gel filtration column running in GF150 buffer on AKTA Purifier FPLC instrumentation. Fractions containing dynactin were pooled and concentrated as above to a final concentration of 1–2 mg/mL, as determined by the Bradford assay.

All purified proteins were dispensed into single-use aliquots, flash frozen in liquid nitrogen, and stored at - 80 °C.

### RNA templates, synthesis, and purification

RNA constructs used for cryo-EM and for MST were synthesized commercially (Horizon Discovery; see Table S4) with the exception of the 151-nucleotide RNA containing the *hairy* localization element, which was transcribed in vitro from a *hairy* 3′ UTR template (*18*) (GenBank accession number X15905), as described below. The *TLS* and *hSL1* constructs (as well as their mutants) included 5′ and 3′ flanks of 8 nucleotides (*TLS*) or 10 nucleotides (*hSL1*) of native sequence from the *K10* and *hairy* 3′ UTRs. RNAs used for MST included a 5′ Cy5 label that was linked to stem loops via a linker of two adenosines (with the exception of the *TLS* and *hSL1* constructs whose native flanking sequence at the 5’ end of the stem loops served as the linker). For single-molecule motility assays with the *TLS* and *hSL1*, Cy3-labeled versions of these transcripts were used in addition to the Cy5-labeled constructs.

*K10* 3′ UTR constructs for single-molecule assays and embryo injection were transcribed from, and numbered according to, the *K10* cDNA (GenBank accession number AY060415). The template for the 1490-nucleotide 3′ UTR construct (positions 1608–3061 of the cDNA) contained the poly-A signal from the cDNA and an additional 38 base pairs that included an SP6 promoter from the backbone that was not counted in the numbering. Mutations of the *TLS* within the *K10* 3′ UTR were introduced using a construct with engineered HindIII and NheI sites on either side of the *TLS* that was previously shown not to affect localization of mRNA following injection into embryos (*52*). Synthetic DNA oligos (Merck) encoding the desired *TLS* mutations and complementarity to the HindIII and NheI overhangs were ligated to the digested plasmid containing the *K10* 3′ UTR, thereby replacing the encoded *TLS* with the mutant sequence. The sequence for the scrambled *TLS*, which is a non-localizing control incapable of making a stem-loop structure, was described previously (*52*). The *hairy* 3′ UTR corresponds to positions 1185–1845 of GenBank accession number X15905 and is part of a 730-nucleotide transcript used for injection. Templates of a mutant version of the *hairy* 3′ UTR in which *hSL1* lacks bulges were synthesized commercially (GenScript). A 573-nucleotide fragment of the *I-factor* RNA coding sequence containing the wild-type *ILS* or a version lacking bulges were transcribed from commercially synthesized template (IDT) and correspond to positions 2932–3498 of GenBank accession number M14954.2. The 839-nucleotide *bicoid* 3′ UTR RNA containing wild-type *bcdSLV* or a version lacking bulges were transcribed from a commercially synthesized template (GenScript) and correspond to positions 1702–2536 of GenBank accession number NM_169159.4. Full length 1,718-nucleotide *gurken* transcripts containing the wild-type *GLS* or a version lacking bulges were transcribed from commercially synthesized templates (GenScript) and correspond to positions 1–1718 of GenBank accession number NM_057220.3.

*LacZ* RNAs with individual *TLS* and *KSE* structures, or a combination of both, were transcribed from a fragment of the *LacZ* gene that was comparable in length to the *K10* 3′ UTR and predicted to code for an RNA that has minimal secondary structure as determined by the greatest average ss-count in mfold predictions (*72*) (https://www.unafold.org/mfold/applications/rna-folding-form.php). At a position analogous to the location of the *TLS* in the *K10* 3′ UTR (673 base pairs), the *TLS* and *KSE* sequences were added to the design and commercially synthesized (GenScript). For the transcript containing both the *TLS* and *KSE*, the linker between the stem loops is the same as the native sequence between *hSL1* and *hSL2 (18)*. This template was also used to transcribe the 145-nucleotide *TLS-KSE* construct for single-molecule motility assays.

Linear DNA templates with a T7 or T3 polymerase promoter were prepared by PCR or by restriction digestion at the 3′ end of the desired transcript. Templates were then agarose gel-purified. RNA was synthesized using MEGAscript T7 (full length *K10*; ThermoFisher) and MEGAshortscript T7 (*hSL1-hSL2* and *TLS-KSE*; ThermoFisher) in vitro transcription kits, or the mMESSAGE mMACHINE T7 or T3 in vitro transcription kit (ThermoFisher) to make 5′-capped transcripts for injection into *Drosophila* embryos. Fluorescent labeling of in vitro transcribed RNA was achieved by stochastic incorporation of Cy3- or Cy5-labeled UTP included in synthesis reactions at a ratio in the reaction mix of 1:4 labeled:unlabeled UTP for single-molecule motility assays or 1:9 labeled:unlabeled UTP for injection assays. Following in vitro synthesis for 2–4 hours at 37 °C, template DNA was digested with DNaseI, and RNA was purified by a single phenol:chloroform:isoamyl alcohol extraction (25:24:1; Ambion) followed by successive passage through two Microspin G50 columns (Cytiva) to remove unincorporated nucleotides. RNA was then precipitated with ammonium acetate/ethanol. RNA concentration was determined using a NanoDrop One spectrophotometer (ThermoFisher) and RNA integrity was confirmed by agarose gel electrophoresis before the RNA was dispensed into aliquots and stored at -80 °C.

### Cryo-EM sample preparation

For assembling Egl–BicD complexes with the *TLS, hSL1, ILS* and *GLS*, 1.5 μM of the full-length Egl–BicD complex was incubated with 25 μM RNA for 45 minutes at 4 °C in GF150 buffer containing 0.00125% IGEPAL (MilliporeSigma). For the Egl–BicD–*bcdSLV* and Egl– BicD–*hSL1–hSL2* complexes, a truncation of BicD (residues 322–782) was used that excludes CC1. For Egl–BicD–*bcdSLV*, 0.75 μM of Egl–BicD(322-782) was incubated with 25 μM of *bcdSLV* RNA. For the Egl–BicD– *hSL1–hSL2* complex, 0.75 μM of Egl–BicD(322-782) was incubated with 2 μM of *hSL1-hSL2* RNA. The sample was then centrifuged at 12,000 x g for 2 minutes. 3.5 μl of the supernatant was applied to freshly glow-discharged Quantifoil R2/2 300-square-mesh gold grids (Quantifoil) in a Vitrobot IV (ThermoFisher) at 95% humidity and 4 °C, incubated for 10 seconds, and blotted for 2 seconds before being plunged into liquid ethane.

### Cryo-EM data collection and image processing

The cryo-EM samples were imaged using a FEI Titan Krios (300 kV) equipped with a K3 detector and an energy filter with a 20 eV slit size (Gatan). Automated data collection was performed using ThermoFisher EPU. All data collection statistics can be found in Table S1.

Egl–BicD–*TLS* complex: 8,568 movies were acquired at a magnification of 81,000x (1.059 Å/pixel) using a 100 μm objective aperture, with 40 frames per movie and a total fluence of ∼48 e⁻/Å². Global motion correction and dose-weighting were performed in RELION-4.0 (*73*) using MotionCorr2 (*74*) with a B-factor of 150 and 5x5 patches. Patch-based CTF estimation and initial processing steps were conducted in CryoSPARC (*75*). Particles were initially picked using an ellipse (180 x 100 Å) as a reference. Subsequent 2D classification was used to identify protein-like densities, which were employed as references for an additional round of reference-based particle picking (Fig. S2). Approximately 5 million particles were extracted with a box size of 200 pixels and a pixel size of 2.12 Å. 2D classification was then performed to select ∼1.6 million particles belonging to classes with protein-like features. Ab-initio reconstruction generated four 3D references, which were subsequently used for heterogeneous refinement. Two of these classes (Class 1 and Class 2; Fig. S2) displayed density corresponding to a coiled-coil region, RNA, and distinct domains of Egl. Particles from these classes were selected for further processing in RELION-4.0 and 5.0 (*76*). A third class (Class 3) was observed which lacked density for ExoHD and XAD. In this class, the RNA and its bound ED1 and ED2 exhibited significant flexibility, which precluded high-resolution 3D refinement.

Approximately 1 million particles from Classes 1 and 2 of the heterogeneous refinement were re-extracted in RELION-4.0 with a box size of 280 pixels and a pixel size of 1.059 Å. This was followed by 3D refinement using Class 2 as the reference. This choice was informed by the observation that the RNA and associated Egl domains on either side of the BicD coiled coil exhibited flexibility relative to each other. To improve particle alignment, one of the ExoHD–XAD modules was omitted from the 3D reference and the mask. Local particle motion was subsequently corrected using particle polishing, during which the particles were re-extracted with the 360-pixel box at 1.059 Å pixel size. Another round of 3D refinement was performed, followed by CTF refinement to refine per-particle defocus, per-micrograph astigmatism, and to estimate beam tilt and trefoil parameters. A final 3D refinement was then conducted, resulting in a 3.1 Å resolution consensus structure. To sort for conformational and compositional heterogeneity, 3D classification without alignment was performed, focusing on either side of the BicD coiled coil. Classification with a mask around the ExoHD–XAD module included in previous 3D refinements enabled sorting of conformations within the stable part of the complex. Two major conformations (Structures A and B; Fig. S2) were identified, differing slightly in the position of the *TLS* RNA and its bound ED2, ExoHD and XAD. Structure A, at the 3 Å resolution, was used for analyzing interactions among the different components of the complex and for identifying sites for mutagenesis as it was the best resolved structure.

In addition to focused 3D classification and refinement, we performed 3D classification with a mask around the full complex that also incorporated the ExoHD–XAD module excluded in prior 3D refinements. This facilitated sorting of Egl–BicD–*TLS* structures with either one or two ExoHD–XAD modules bound, as well as the conformations within these particle populations.

Approximately half of the particles contained two ExoHD–XAD modules, while the other half had only one. Among the structures with two ExoHD–XAD modules, distinct conformations (Structures C, D, and E; Fig. S2) were observed. These conformations shared consistent RNA-binding interactions across the domains but differed in the relative orientation of the RNA and its associated domains with respect to the BicD coiled coil. Structure C, at 3.4 Å resolution, was the best resolved structure with all domains engaged with the RNA and was therefore used to describe the architecture of the complex in Fig. 1.

Egl–BicD–*hSL1* complex: one dataset with 12,420 movies was acquired with 1.06 Å/pixel, 50 frames per movie and a total fluence of ∼50 e⁻/Å², and another with 16,453 movies acquired with 0.91 Å/pixel, 56 frames per movie and a total fluence of ∼52 e⁻/Å². For each dataset, global motion correction and dose-weighting were performed in RELION-4.0 using MotionCorr2 with a B-factor of 150 and 5x5 patches. Patch-based CTF estimation and initial processing steps were conducted in CryoSPARC. Particles were picked using an ellipse (180 x 100 Å) as a reference and approximately 9.1 million particles were extracted in total at a box size of 90 pixels and binning factor of four. 2D classification was then used to select approximately 4.9 million particles that belonged to 2D classes displaying protein-like densities (Fig. S6). Ab-initio reconstruction was used to generate four 3D references, which were subsequently used for two rounds of heterogeneous refinement. The class displaying density corresponding to a coiled-coil region, RNA, and distinct domains of Egl (Class 1; Fig. S6) was selected after each round of heterogeneous refinement. The selected particles were re-extracted at a box size of 320 pixels (1.06 Å/pixel) or 360 pixels (0.91 Å/pixel), followed by another round of ab-initio reconstruction. A subset of 560,115 particles, showing well-defined densities for Egl–BicD (Class 1 and 2; Fig. S6), were selected for further processing in RELION-4.0 and 5.0.

Particles from both the datasets were merged at this stage and a round of 3D refinement was performed. Local particle motion was subsequently corrected using particle polishing. Another round of 3D refinement was performed, followed by CTF refinement to refine per-particle defocus, per-micrograph astigmatism, and to estimate beam tilt and anisotropic magnification parameters. A final 3D refinement was then conducted, resulting in a 3.3 Å resolution consensus structure. To sort for conformational and compositional heterogeneity, 3D classification without alignment was performed, focusing on either side of the BicD coiled coil. Classification with a mask around the ExoHD–XAD module included in previous 3D refinements enabled sorting of conformations within the stable part of the complex. One major conformation (Structure A; Fig. S6) was identified for this part of the complex and was resolved at an overall resolution of 3.2 Å. Additionally, classification with a mask around the ExoHD–XAD module excluded in prior 3D refinements facilitated sorting of Egl–BicD–*hSL1* structures with either one or two ExoHD–XAD modules bound, as well as the conformations within these particle populations. Approximately half of the particles contained two ExoHD–XAD modules, while the other half had only one. Among the structures with two ExoHD–XAD modules, distinct conformations (Structures B and C; Fig. S6) were resolved at an overall resolution of 3.9–4.1 Å. These conformations shared consistent RNA-binding interactions across the domains but differed in the relative orientation of the RNA and its associated domains with respect to the BicD coiled coil. Structure A resolved at the highest resolution and was therefore used for analyzing interactions.

Egl–BicD–*ILS* complex: 8,009 movies were acquired at a magnification of 81,000x (1.059 Å/pixel) using a 100 μm objective aperture, with 40 frames per movie and a total fluence of ∼47 e⁻/Å². Patch motion correction and Patch-based CTF estimation were performed in CryoSPARC. Particles were picked using an ellipse (180 x 100 Å) as a reference and approximately 4.4 million particles were extracted at a box size of 200 pixels and a pixel size of 4.24 Å. 2D classification was then used to select approximately 2 million particles that belonged to 2D classes displaying protein-like densities (Fig. S7). Ab-initio reconstruction was used to generate three 3D references, which were subsequently used for heterogeneous refinement. The class displaying density corresponding to a coiled-coil region, RNA, and distinct domains of Egl (Class 1; Fig. S7) was selected for another round of ab-initio followed by heterogeneous refinement. A subset of 360,597 particles that had well-defined densities for Egl–BicD (Class 1; Fig. S7) was selected for further processing in RELION-5.0. Local particle motion was subsequently corrected using particle polishing, during which the particles were re-extracted with the 360-pixel box at 1.059 Å pixel size. 3D refinement was then performed using a mask that excluded one of the ExoHD–XAD modules, focusing on the most stable regions of the complex. This yielded a 3.4 Å resolution structure of the Egl–BicD–*ILS* complex (Structure A; Fig. S7).

To sort for compositional heterogeneity of the ExoHD– XAD module, heterogeneous refinement was performed, classifying particles into two groups: one with a single ExoHD–XAD module and another with two. Approximately 57% of particles contained one bound ExoHD–XAD module, while 43% contained two. The 3D refinement of the class with two ExoHD–XAD modules resulted in a 3.7 Å resolution structure (Structure B; Fig. S7). However, local resolution distribution indicated that the ExoHD–XAD modules exhibited flexibility relative to each other, leading to decreased local resolution in these regions when both modules were included in the mask for refinement. Due to its higher quality, the 3.4 Å resolution structure was used for model building and to elucidate the binding mechanism of Egl to the *ILS*.

Egl–BicD–*bcdSLV* complex: 22,750 movies were acquired at a magnification of 81,000x (1.059 Å/pixel) using a 100 μm objective aperture, with 50 frames per movie and a total fluence of ∼50 e⁻/Å². Patch motion correction and Patch-based CTF estimation were performed in CryoSPARC. Approximately 13 million particles (200-pixel box; 2.12 Å/pixel) were initially picked using an ellipse (180 x 100 Å) as a reference (Fig. S8). In parallel, 3.3 million particles were also picked using Topaz (*77*) with a model trained using a small subset of 12,000 particles selected from 2D classification. Subsequently, a round of heterogeneous refinement using 3D references similar to those used for the *TLS*-bound structure was performed. Classes showing defined features for Egl–BicD–RNA were selected for a round of 3D classification without alignment. The particles coming from template-based picking and Topaz picking were then merged. After removal of duplicate particles, approximately 880,000 particles remained. Local particle motion was subsequently corrected in RELION-5.0, during which the particles were re-extracted with a 360-pixel box at 1.059 Å pixel size. Another round of 3D refinement was performed, resulting in a 3.3 Å resolution consensus structure. To sort for conformational and compositional heterogeneity, 3D classification without alignment was performed, focusing on either side of the BicD coiled coil. Classification with a mask around the ExoHD–XAD module included in previous 3D refinements enabled sorting of conformations within the stable part of the complex. One major conformation (Structure A; Fig. S8) was identified for this part of the complex after 3D classification. The selected particles were then used for 3D refinement followed by CTF refinement (Beam tilt and Trefoil parameters) and finally another round of 3D refinement, which resulted in a 3.4 Å resolution structure. Additionally, classification with a mask around the ExoHD–XAD module excluded in prior 3D refinements facilitated sorting of Egl–BicD–*bcdSLV* structures with either one or two ExoHD–XAD modules bound, as well as the conformations within these particle populations. Approximately 37% of the particles contained two ExoHD–XAD modules, while 53% had only one. Among the structures with two ExoHD–XAD modules, distinct conformations (Structures B and C; Fig. S8) were sorted after an additional round of 3D classification with the mask around the whole molecule and resolved at an overall resolution of 4.2–4.4 Å. However, local resolution distribution indicated that the ExoHD–XAD modules exhibited flexibility relative to each other, leading to decreased local resolution in these regions when both modules were included in the mask for refinement. Structure A resolved at the highest resolution and was therefore used for analyzing interactions.

Egl–BicD–*GLS* complex: 12,592 movies were acquired at a magnification of 81,000x (0.91 Å/pixel) using a 100 μm objective aperture, with 56 frames per movie and a total fluence of ∼53 e⁻/Å². Global motion correction and dose-weighting were performed in RELION-5.0 using MotionCorr2 with a B-factor of 150 and 5x5 patches. Patch-based CTF estimation and initial processing steps were conducted in CryoSPARC. Approximately 6 million particles were initially picked using an ellipse (180 x 100 Å) as a reference (Fig. S9). Subsequently, a round of heterogeneous refinement using 3D references similar to those used for the *TLS*-bound structure was performed (Fig. S2). A class showing defined features for Egl–BicD–RNA was selected for a round of 3D classification without alignment. The best-defined 3D class was selected and two more iterations of heterogeneous refinement and 3D classification were performed while selecting the 3D class showing defined features for Egl–BicD–RNA at each step. This resulted in a total of 41,216 particles which were then used to train a picking model in Topaz. The picked particles (∼ 2.3 million) were then sorted using three rounds of heterogeneous refinement and 3D classification as was done for particles picked using an elliptical reference resulting in a set of 66,615 particles. Both sets of selected particles were merged followed by removal of duplicated particles. For this complex, while there were classes from 2D classification that showed two ExoHD–XAD modules engaged with the complex, the low number of overall particles hindered the classification of complexes with one or two ExoHD–XAD modules in 3D. Therefore, after initial particle sorting the processing was focused on resolving the complex while having only one ExoHD–XAD module within the mask. 3D refinement in RELION-5.0 was subsequently performed followed by particle polishing and CTF refinement (per particle defocus and per micrograph astigmatism), resulting in a 3.9 Å resolution structure which was used to analyze interactions.

Egl–BicD–*hSL1*–*hSL2* complex: 44,979 movies were acquired at a magnification of 81,000x (1.059 Å/pixel) using a 100 μm objective aperture, with 50 frames per movie and a total fluence of ∼50 e⁻/Å². Initial image processing steps including particle picking and sorting were performed as described for the Egl–BicD–*bcdSLV* structure (Fig. S8). A total of 1.14 million particles were obtained after initial particle sorting (Fig. S15). Local particle motion was subsequently corrected in RELION-5.0, during which the particles were re-extracted with the 360-pixel box at 1.059 Å pixel size. Another round of 3D refinement was performed resulting in a consensus structure. To sort for compositional heterogeneity, 3D classification without alignment was performed using a mask encompassing the full complex. We found that ∼60% (675,190) of the particles had Egl–BicD bound to both *hSL1* and *hSL2*, representing the Egl–BicD–*hSL1– hSL2* complex. The remaining particles displayed both stem-loop binding sites occupied by *hSL1* stem loops, resembling the Egl–BicD–*hSL1* complex.

To sort for conformational heterogeneity within Egl– BicD–*hSL1–hSL2* complex, another round of 3D classification without alignment was performed using a mask encompassing the full complex. This identified a major conformation, which was used for further 3D refinement, yielding a map at 3.4 Å resolution. This structure was used to analyse the interaction of *hSL2* with Egl–BicD.

Although the linker between *hSL1* and *hSL2* in the Egl– BicD–*hSL1*–*hSL2* structure appeared flexible, we detected additional low-resolution density connecting the two stem loops (Fig. S15C) which was absent in the structures with only *hSL1* bound. This observation therefore shows that both stem loops in the Egl–BicD– *hSL1*–*hSL2* structure originate from the same RNA molecule (Fig. S15C).

Notably, we did not observe 3D classes where the ExoHD–XAD module was bound to the *hSL2* stem loop; in contrast, this module was consistently bound to only *hSL1*. Further, in cases where both stem-loop binding sites were occupied by *hSL1* either in this dataset or in the Egl–BicD–*hSL1* complex dataset, we observed structural classes with two ExoHD–XAD modules bound, in addition to those with only one (Fig. S6 and S15). Thus, our data together with structural observations presented in Fig. 4 suggests that *hSL2* is not compatible with ExoHD–XAD binding.

### Model building and refinement

For modeling Egl, the AlphaFold2-generated (*38*) model (AF-Q9W1K4-F1) of *Drosophila melanogaster* Egl isoform B (Uniprot Q9W1K4) was obtained from the AlphaFold Protein Structure Database (*78*). The individual domains (ED1, ED2, ExoHD and XAD) were fit into the cryo-EM density using UCSF ChimeraX (*79*) guided by the side chain densities and the shape of the domains. Given the flexible linkers between domains in Egl, their assignment to specific Egl chains was guided by multiple structural criteria. ED1 and ED2, positioned across the BicD coiled coil, were assigned to the same Egl molecule based on a cryo-EM density at low threshold connecting these domains and a linker length compatible with this arrangement. The ExoHD–XAD module was assigned to the same chain as it is adjacent to ED2. This was supported by an interaction between residues 386–394 and the ExoHD, which reduces the intervening sequence between ED2 and ExoHD to approximately 217 residues. Additional folded elements within this region likely shorten the effective linker length even further, making it more likely that the ExoHD adjacent to ED2 belongs to the same chain. Although less favorable, it is possible that some ExoHD–XAD module might come from a different Egl molecule to their neighboring ED2.

For modeling BicD, an AlphaFold2 prediction of *Drosophila melanogaster* BicD (Uniprot P16568) was used. The model was fit into the cryo-EM map using UCSF ChimeraX guided by the side chain densities and an AlphaFold2 prediction of ED1 (residues 1–82) bound to BicD residues 700–782.

For modeling the RNA stem loops, 3D structures predicted by Alphafold3 server (*80*) or those generated by RNAComposer server (*81*) were used as starting models.

After placing individual protein domains or RNA stem loops into the cryo-EM map, the model went through iterative cycles of restrained flexible fitting using ISOLDE (*82*), followed by user-guided refinement in ISOLDE or COOT (*83*). Final model refinement and model validation was performed in PHENIX (*84*). All refinement statistics can be found in Table S1.

### AlphaFold2 prediction

All structure predictions unless specified were performed using AlphaFold2 (*38*) (for single chains) or AlphaFold2-Multimer (*85*) (for multiple chains) through a local installation of ColabFold (*86*). Both AlphaFold2 and AlphaFold3 accurately predicted the interaction between ED1 and BicD. However, we were unable to obtain reliable predictions for the Egl–BicD complex bound to different RNA targets. This highlights the ongoing challenge of predicting protein–RNA complexes and the need for more experimentally-determined structures.

### RNA secondary structure prediction

RNA secondary structures were predicted using the RNAfold web server (*87*): http://rna.tbi.univie.ac.at/cgi-bin/RNAWebSuite/RNAfold.cgi Notably, some nucleotides predicted to be part of bulges or loops were found to be integrated into the dsRNA helix in the experimental structures, where they participated in stacking and non-canonical base pair interactions (Fig. S10B). Therefore, when generating the Δb constructs, nucleotide bulges were removed based on experimental structures whenever discrepancies arose between the predicted and experimental models.

### Microscale thermophoresis

The affinity of the Egl–BicD complex and its variants for the indicated RNA stem loops was measured using a NanoTemper Monolith instrument and MO.Control MST acquisition software (NanoTemper). Two-fold serial dilutions of the Egl–BicD complex ranging from 1.59 x 10^-10^ M to 7.70 x 10^-6^ M (assuming 2:2 stoichiometry of Egl:BicD) were incubated with 1 nM Cy5-labeled RNA at room temperature for 15 minutes in GF150 buffer supplemented with 0.05% Tween-20. Higher concentrations of Egl–BicD could not be tested as they were prone to aggregation. Serial dilutions were loaded into standard capillaries (MO-K022) and irradiated with infrared light at room temperature for 10 seconds (medium MST power) at 20% excitation, with changes in fluorescence monitored by the pico-red detector. MST traces were analyzed in MO.Affinity Analysis with data fitting performed using a 1.5 second on-time (F_hot_ = 0.5– 1.5 seconds; F_cold_ = -0.5–0 seconds) and the *K*_d_ model, which was able to fit binding curves in the absence of saturated RNA binding given sufficient signal to noise (minimum value of 5.0 to be considered interacting). Data were further analyzed in GraphPad Prism (Version 10.4.0) to determine the standard deviation (SD) of the fit values of *K*_d_ and to enable statistical comparison of different conditions.

### Injection of *Drosophila* embryos with fluorescent RNA

Wild-type embryos (*w^1118^* strain) were collected and injected with Cy3-labeled RNA as described previously (*18, 46, 52*). For a typical experiment, up to 60 dechorionated embryos were mounted in Voltalef oil 10S for injection of syncytial blastoderms with a 250 ng/µl solution of RNA. The person performing the injections was blinded to the identity of the RNA being evaluated. Following the last injection, embryos were incubated at room temperature for 8 minutes (∼13 minutes from injection of the first embryo) before fixation with formaldehyde-saturated heptane, manual removal of the vitelline membrane with fine syringe needles and mounting in VECTASHIELD antifade medium with DAPI (Vector Labs) for visualization of nuclei.

Imaging was conducted using a Zeiss LSM 710 or 780 confocal microscope equipped with a 40x oil-immersion objective (1.3 NA). A zoom factor of 1.8 was used to capture images with 684 x 512 pixels and the laser intensity was adjusted to allow visualization of injected RNA without reaching saturation. The position for image capture was determined by selecting the Z-plane in which both fluorescently-labeled RNA and the longest axis of the nuclei were visible.

The efficiency of RNA localization in injected embryos was quantified by comparing in FIJI (*88*) the apical and basal RNA intensity within uniform regions of interest (ROIs), whose size (7.391 µm^2^) was predefined as the mean area occupied by apically-localized RNA at individual microtubule organizing centers (positioned just above nuclei) in ∼100 embryos injected with the wild-type *K10* 3′ UTR. For each image, four ROIs were placed in the apical region of the injection site, with each ROI centered on the brightest area of fluorescence intensity. Corresponding basal ROIs were translated vertically to a position just above the yolk. The mean fluorescence intensities of these ROIs were then averaged and background-corrected by subtracting the mean intensity of a basal region distant from the injection site before being expressed as a ratio of apical to basal intensity. This analysis was applied to 22–77 blastoderm embryos from 2–6 independent injections per condition.

### Single-molecule-resolution RNA motility assays

Total internal reflection fluorescence (TIRF)-based motility assays of reconstituted dynein transport complexes assembled with the indicated RNAs were performed as previously described (*34*). Briefly, assembly mixes of 100 nM dynein, 500 nM Egl–BicD, 200 nM dynactin, and 50 nM RNA (25 nM each of Cy3- and Cy5-labeled samples) were incubated in a total volume of 5 µl GF150 on ice for 1 hour prior to imaging. Complexes were assembled with a 10-fold molar deficit of total RNA relative to Egl–BicD to assess the sufficiency of single RNAs to activate motility, which may be obscured by excess RNA that drives Egl–BicD occupancy beyond that needed to initiate transport. The assembly mixes were diluted 10- to 80-fold in motility buffer (30 mM HEPES pH 7.3, 50 mM KCl, 5 mM MgSO_4_, 1 mM EGTA pH 7.5, 1 mM DTT, 20 µM Taxol (Sigma), 0.5 mg/mL BSA, 1 mg/mL alpha-casein (Sigma)) with 1 mM MgATP and an oxygen scavenging system (1.25 μM glucose oxidase, 140 nM catalase, 71 mM 2-mercaptoethanol, 25 mM glucose) and applied to an ∼10-µl flow chamber containing streptavidin-immobilized microtubules (labeled with HiLyte 488- and biotin-porcine tubulin; Cytoskeleton Inc) on a PEG-biotin-passivated cover slip. Motility of Cy3- and Cy5-labeled RNA within these chambers was alternately recorded with an iXon^EM^+ DU-897E EMCCD camera (Andor) mounted on a Nikon TIRF system with a Nikon APO TIRF 100x oil objective (1.49 NA) using Micro-manager acquisition software (*89*) at the maximum possible framerate (∼2 frames per second for each channel) and 100 ms exposure for each channel. Samples were illuminated with a 150 mW Coherent Sapphire 488 nm laser for HiLyte 488-labeled microtubules, a 150 mW Coherent Sapphire 561 nm laser for Cy3-labeled RNA, and a 100 mW Coherent CUBE 641 nm laser for Cy5-labeled RNA. Assemblies of dynein–dynactin–BicD–Egl complexes with Cy3-and Cy5-labeled *TLS*-*KSE* constructs were imaged using a Nikon Ring-TIRF system controlled by Micro-manager and equipped with the iLas 2 platform (GATACA Systems) and the same objective as above. These samples were illuminated for 200 ms with 488 nm (for HiLyte 488-labeled microtubules), 561 nm (for Cy3-RNA), and 647 nm (for Cy5 RNA) lasers within a Cairn Multiline Kompact laser box and the motility in Cy3 and Cy5 channels recorded simultaneously on independent Photometrics Prime 95B CMOS cameras at the maximal frame rate (∼4 frames per second).

Colocalization of Cy3 and Cy5 RNA was analyzed manually in FIJI using kymographs derived from acquisitions described above.

### RNA bend analysis

A custom Python program was used to calculate the bend between the upper and lower helices of RNA localization signals. First, the Curves+ software (*90*) was used to obtain a helical trajectory for each RNA stem-loop. From these trajectories, the coordinates representing the upper and lower helical regions were separately selected and fitted to best-fit lines in 3D space using a singular value decomposition (SVD) algorithm. Each best-fit line was defined by a centroid and a principal axis vector, and the bend angle (in degrees) was determined by computing the dot product of the two principal axis vectors and then applying the inverse cosine.

### Data visualization and statistical analysis

Cryo-EM maps and models were rendered using ChimeraX. Particle angular distribution was plotted using starparser (*91*). Background-subtracted kymographs were generated using FIJI. Images of *Drosophila* embryos injected with fluorescent RNA were analyzed using unsaturated raw pixel values; however, to aid in visualization of apical vs basal RNA localization, lookup tables in representative images had maximum pixel values set to the mean intensity plus 10 standard deviations and the minimum pixel values set to the mode (typically 0 or 1). Plotting of data and statistical analyses were performed as indicated in the figure legends using GraphPad Prism (Version 10.4.0). Unless otherwise stated, the normality of datasets was assumed but not explicitly tested.

## Acknowledgments

We thank members of the Carter and Bullock groups for advice and support. We are grateful to the LMB Electron Microscopy Facility for access to, and support with, electron microscopy sample preparation and data collection. We thank J. Grimmett, T. Darling, and I. Clayson of LMB Scientific Computing for providing resources, the LMB media preparation team for supplying fly food, and S. McLaughlin from the LMB Biophysics Facility for assistance with MST measurements. Work in the Carter and Bullock groups is funded by the Medical Research Council as part of UK Research and Innovation (UKRI) (file reference numbers MC_UP_A025_1011 and MC_U105178790, respectively). The work was also supported by a UKRI Biotechnology and Biological Sciences Research Council (BBSRC) project grant to S.L.B. (BB/T00696X/1), an EMBO Postdoctoral Fellowship to K.S. (ALTF 197-2021), and a UKRI BBSRC PhD studentship to S.C. (Project reference 2273135 as part of BB/M011194/1).

## Author contributions

K.S., M.A.M., A.C. and S.L.B conceived the project. K.S., S.C., M.A.M. and S.L.B. generated data. All authors analyzed data and wrote the manuscript. A.P.C. and S.L.B. supervised the project.

## Competing interests statement

The authors declare no competing interests.

## Data availability

Atomic coordinates and cryo-EM maps have been deposited in the Protein Data Bank (PDB) or Electron Microscopy Data Bank (EMDB), respectively, under accession codes 9RVY and 54292 (Egl–BicD–*TLS* structure A), 9RVZ and 54293 (Egl–BicD–*TLS* structure B), 9RW0 and 54294 (Egl–BicD–*TLS* structure C), 9RW1 and 54295 (Egl–BicD–*TLS* structure D), 9RW2 and 54296 (Egl–BicD–*TLS* structure E), 9RW3 and 54297 (Egl–BicD– *hSL1* structure A), 9RW4 and 54298 (Egl–BicD–*hSL1* structure B), 9RW5 and 54299 (Egl–BicD–*hSL1* structure C), 9RW6 and 54300 (Egl–BicD–*ILS* structure A), 9RW7 and 54302 (Egl–BicD–*bcdSLV* structure A), 9RW8 and 54303 (Egl–BicD–*bcdSLV* structure B), 9RW9 and 54304 (Egl–BicD–*bcdSLV* structure C), 9RWA and 54305 (Egl– BicD–*GLS*), 9RWB and 54306 (Egl–BicD–*hSL1*–*hSL2*). For Egl–BicD–*ILS* structure B, only the map was deposited in EMDB under the accession code 54301. The data will be released upon formal publication of the manuscript.

## Supplementary Figures

**Fig. S1.**
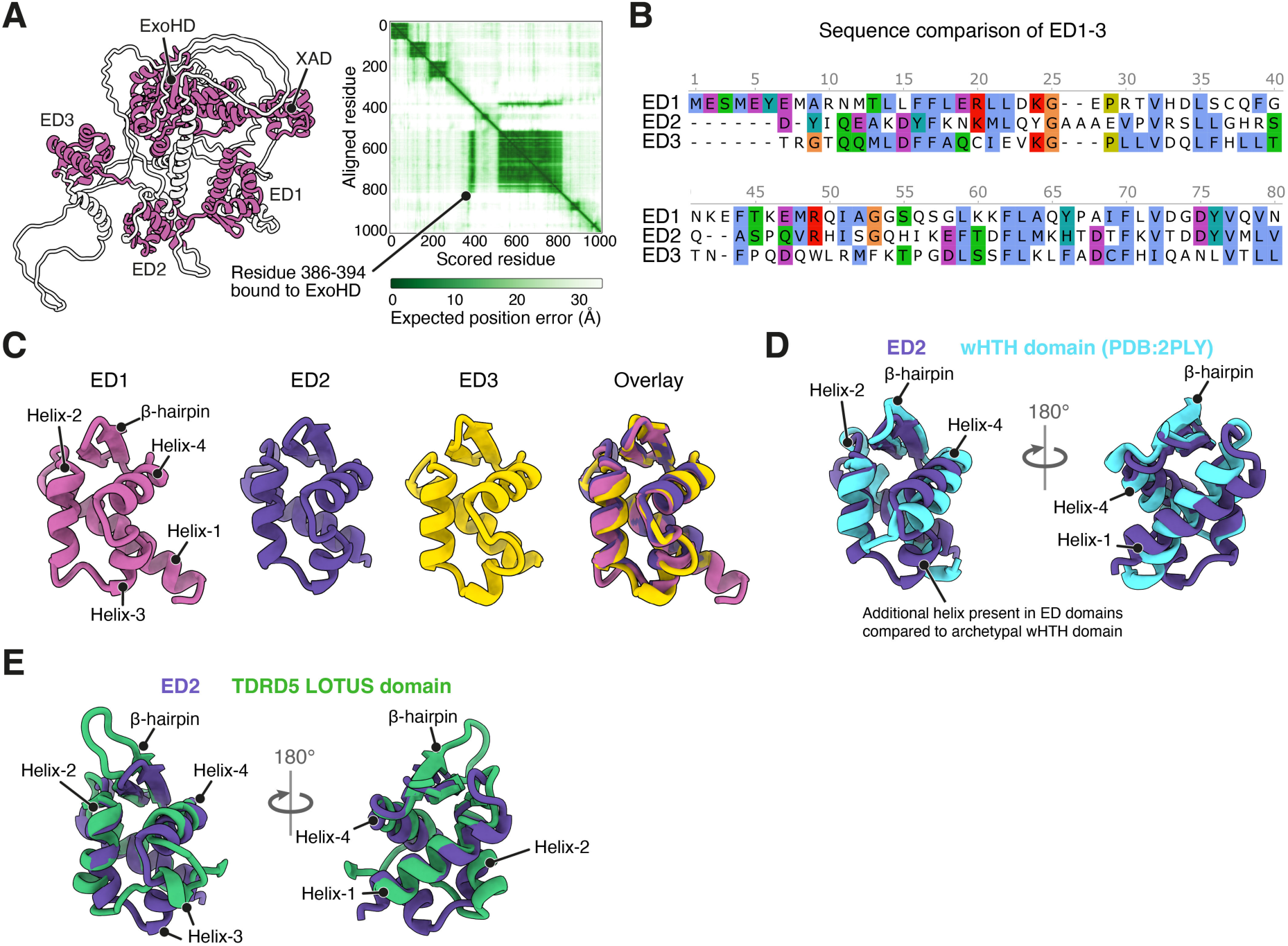
Predicted domains in Egl. (**A**) Alphafold2 prediction of Egl with associated predicted aligned error (PAE) plot (dark green indicates high degree of confidence of local predictions). Structured domains within the RNA-binding region of Egl are shown in magenta. (**B**) Sequence alignments of ED1–3. Alignments are colored according the Clustal X color scheme in which blue represents hydrophobic residues, magenta represents negatively charged residues, green represents polar residues, cyan represents aromatic residues, red represents positively charged residues, orange represents glycines, and yellow represents prolines. (**C**) AlphaFold2 predictions of ED1–3, showing similar winged helix-turn-helix (wHTH) folds. (**D**) Comparison of predicted structures of EDs (using ED2 as representative) with a canonical wHTH fold in the selenocysteine-specific tRNA elongation factor SelB (*92*) showing the addition of ED helix-3 to the canonical wHTH archetype. (**E**) Comparison of predicted structures of EDs (using ED2 as representative) with the Alphafold2 prediction for the LOTUS domain of human TDRD5 (residues 294–396 of Uniprot ID Q8NAT2).

**Fig. S2.**
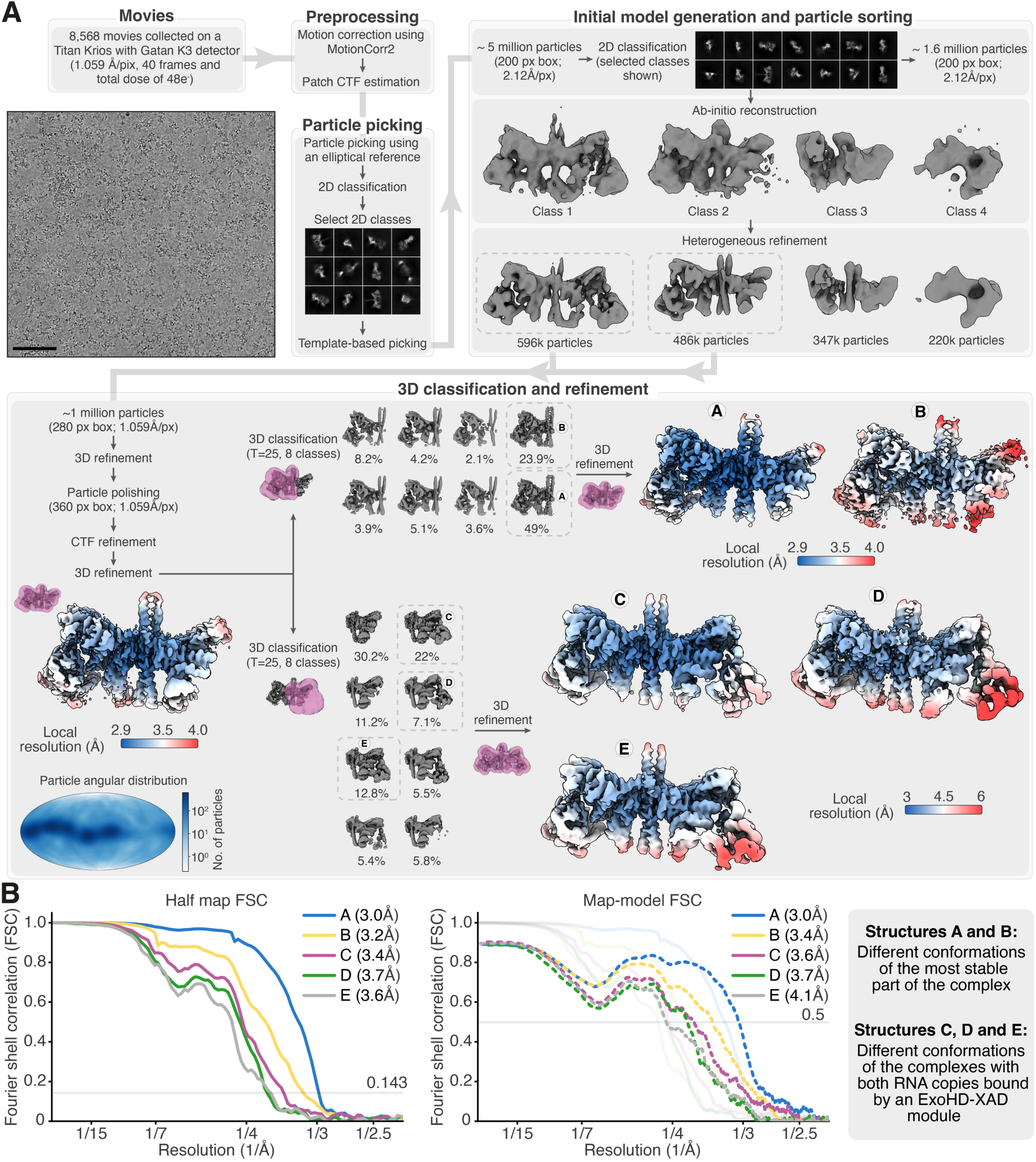
Cryo-EM image processing pipeline for the Egl–BicD–*TLS* complex. (**A**) Image processing was performed using cryoSPARC and RELION-4.0/5.0 and is described in detail in the Materials and Methods section. The classes selected after heterogeneous refinement and 3D classification are indicated with a dotted box. Unless otherwise specified, 3D classifications were performed without alignment (T = Tau fudge). The masks used for 3D classification and 3D refinements are shown in magenta. The angular distribution of particles, projected onto a Mollweide map, shows the orientation coverage of the particles used to obtain the consensus reconstruction. The consensus and final refined maps (structures A–E) are colored based on local resolution estimates from RELION. (**B**) Fourier Shell Correlation (FSC) for structures A–E. The left panel shows the FSC between two independently refined half-maps, with resolution estimated using the 0.143 cutoff. The right panel displays the FSC between the final map and its corresponding model, with resolution estimated using the 0.5 cutoff.

**Fig. S3.**
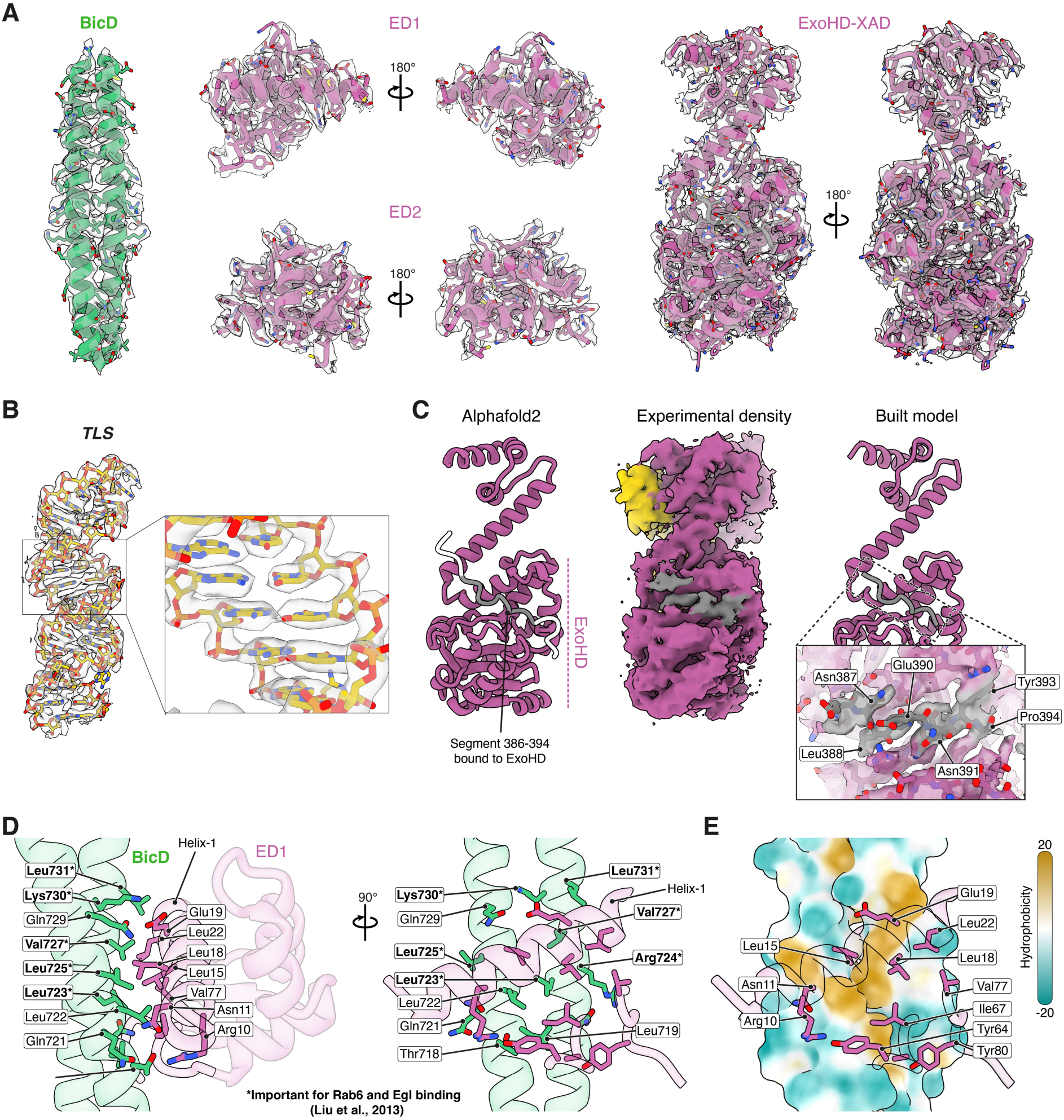
Cryo-EM density and modeled residues in the Egl–BicD–*TLS* complex. (**A** and **B**) Cryo-EM density and modeled residues for Egl–BicD (A) and the *TLS* stem loop (B) within the Egl–BicD– *TLS* structure. (**C**) Alphafold2 prediction of a nine-residue segment (residues 386–394, gray) bound to the ExoHD (left), compared to the experimental density of this region (center) and the model built into the density (right). (**D**) Interaction interface between BicD and ED1, highlighting residues previously shown (*93*) to be important for BicD binding by Rab6 and Egl (bolded). (**E**) Surface representation of the BicD coiled coil, colored by hydrophobicity (brown = hydrophobic, teal = hydrophilic), revealing a hydrophobic patch at the ED1 binding site. A cartoon representation of the interacting ED1 residues is overlaid.

**Fig. S4.**
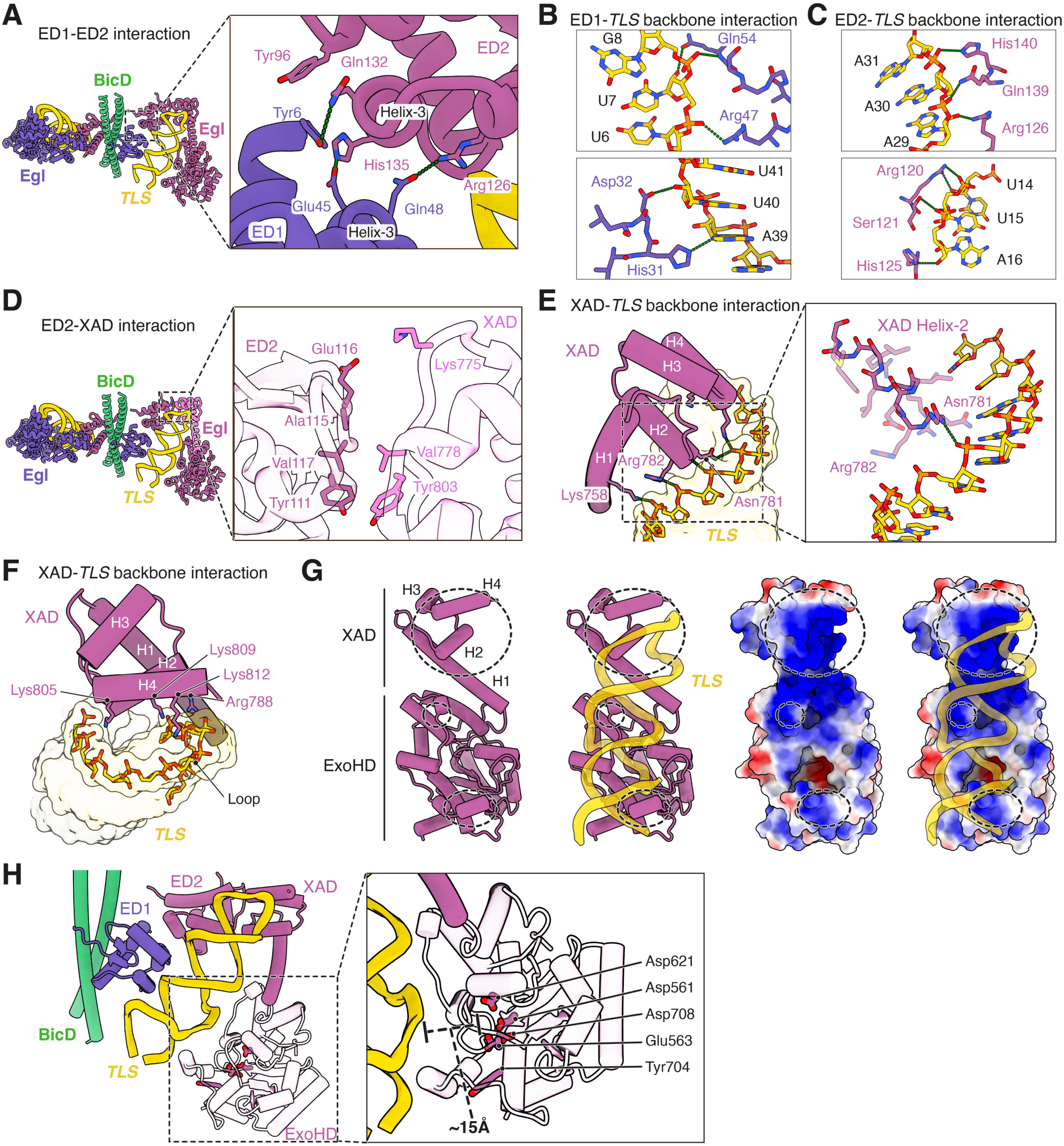
Interactions within the *TLS* binding pocket of Egl. (**A**) Binding interface between ED1 and ED2 when bound to the *TLS.* The interactions are driven by polar interactions which are shown in green. (**B** and **C**) Interactions of ED1 (B) and ED2 (C) with the ribose-phosphate backbone of the *TLS*. (**D**) Binding interface between ED2 and the XAD when bound to the *TLS.* The interactions are primarily hydrophobic in nature driven by Val and Tyr residues. (**E** and **F**) Interactions between the XAD and the ribose-phosphate backbone of the upper helix and loop of the *TLS*. The N-terminal dipole of XAD helix-2 (H2) interacts with the phosphate backbone of the upper helix (E), whereas helices-2 and -4 make contacts across the major groove at the stem-loop junction (F). (**G**) Cartoon representation (left) and surface charge representation (right) of the ExoHD–XAD overlaid with the bound *TLS* stem loop (blue: positive charge, red: negative charge, white: neutral). Positively-charged regions of the ExoHD–XAD that bind the *TLS* are highlighted with a dotted circle. (**H**) Location of the putative catalytic residues of the ExoHD and their average distance from the bound *TLS* stem loop are shown.

**Fig. S5.**
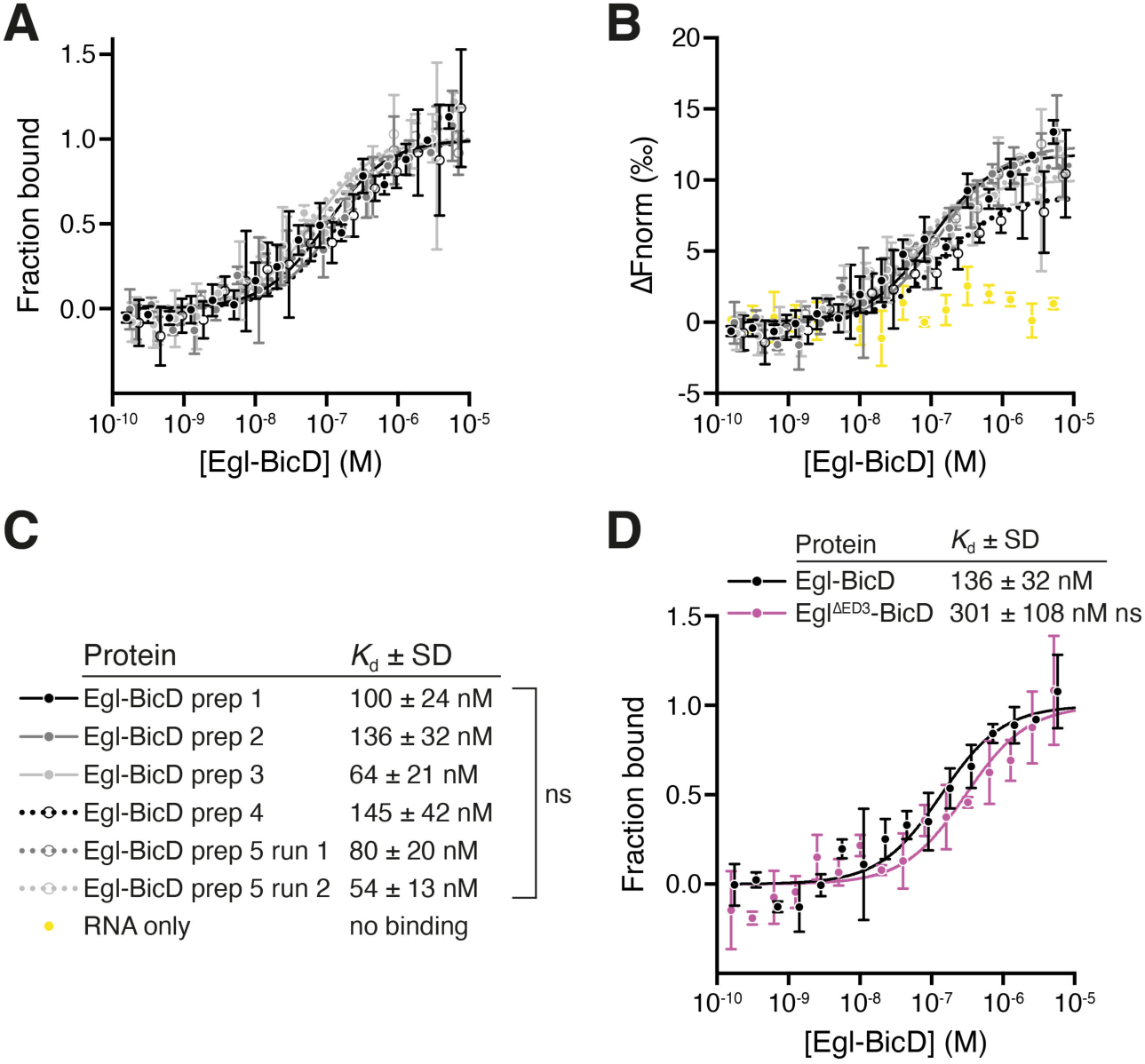
A robust MST assay reveals that ED3 does not contribute significantly to *TLS* binding. (**A** to **C**) MST curves for the *TLS* bound to independent preparations of wild-type Egl–BicD used in this study. Each curve is normalized to show either the fraction of *TLS* that is bound by Egl–BicD (A), or the response amplitude (ΔFnorm (‰); B) relative to a *TLS*-only control (yellow dots). Data points show the mean ± SD for 3–6 replicates for each independent preparation, from which best-fit values for *K*_d_ ± SD were derived (C). No significant deviation in affinity constants is seen between preparations, revealing that this is a robust assay. (**D**) MST binding curve for the *TLS* bound to wild-type Egl–BicD or a variant lacking ED3. Data points show the mean ± SD for 3 replicates per condition, from which best-fit values for *K*_d_ ± SD were derived. Statistical significance was determined using an extra sum-of-squares F test. ns: not significant (p ≥ 0.05). ****p < 0.0001.

**Fig. S6.**
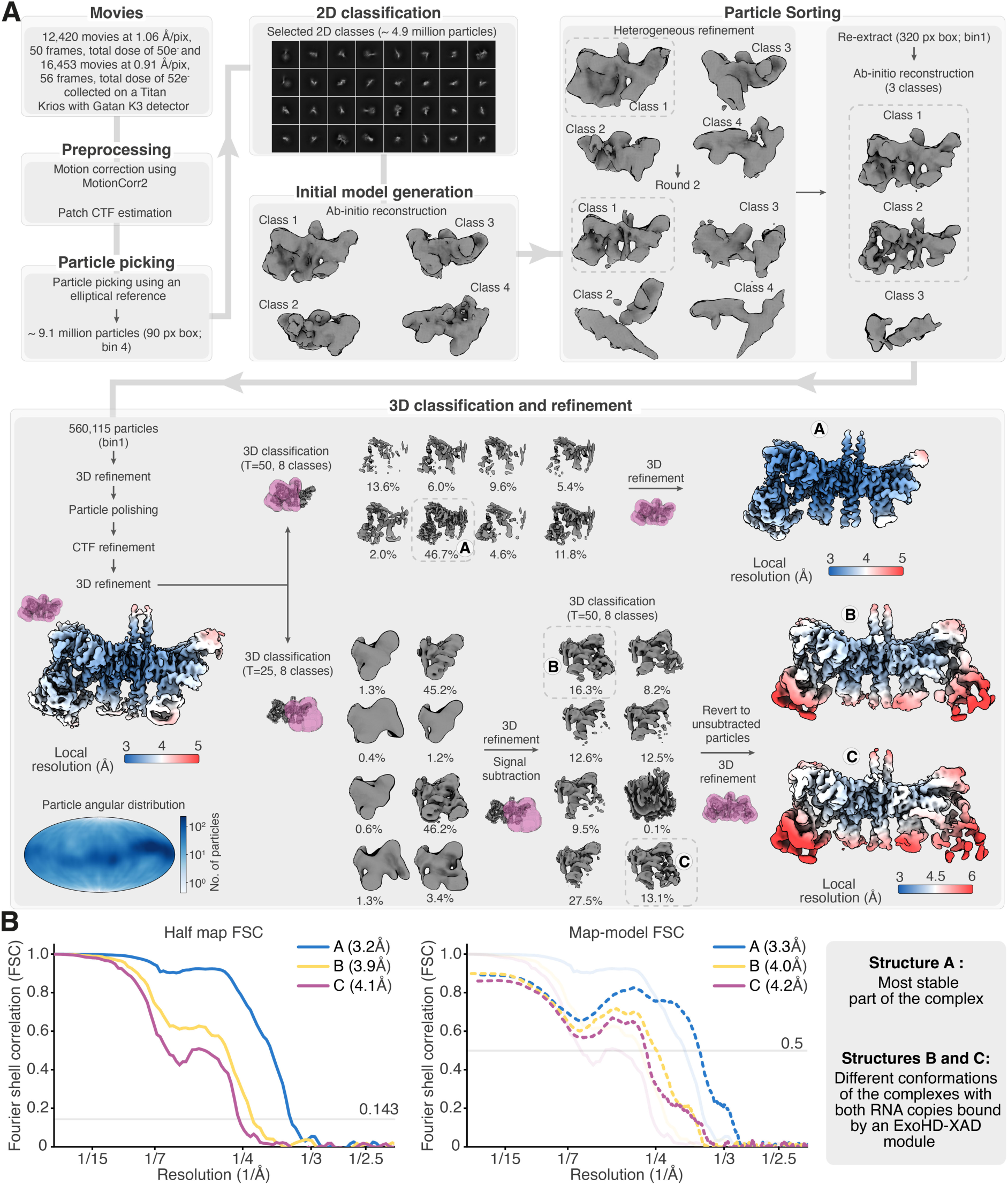
Cryo-EM image processing pipeline for the Egl–BicD–*hSL1* complex. (**A**) Image processing was performed using cryoSPARC and RELION-4.0/5.0 and is described in detail in the Materials and Methods section. The classes selected after heterogeneous refinement and 3D classification are indicated with a dotted box. Unless otherwise specified, 3D classifications were performed without alignment (T = Tau fudge). The masks used for 3D classification and 3D refinements are shown in magenta. The angular distribution of particles, projected onto a Mollweide map, shows the orientation coverage of the particles used to obtain the consensus reconstruction. The consensus and final refined maps (structures A–C) are colored based on local resolution estimates from RELION. (**B**) Fourier Shell Correlation (FSC) for structures A–C. The left panel shows the FSC between two independently refined half-maps, with resolution estimated using the 0.143 cutoff. The right panel displays the FSC between the final map and its corresponding model, with resolution estimated using the 0.5 cutoff.

**Fig. S7.**
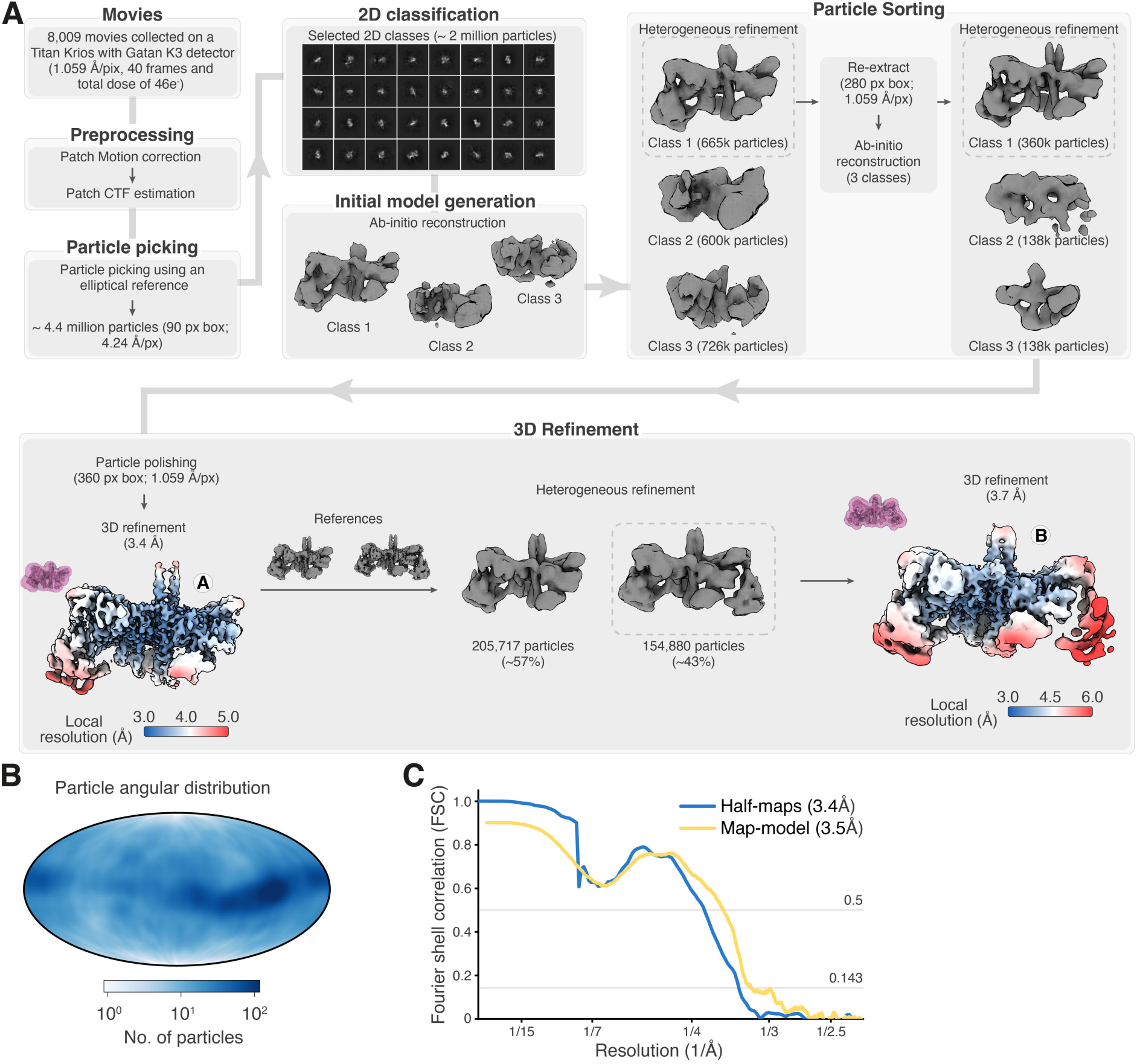
Cryo-EM image processing pipeline for the Egl–BicD–*ILS* complex. (**A**) Image processing was performed using cryoSPARC and RELION-4.0/5.0 and is described in detail in the Materials and Methods section. The classes selected after heterogeneous refinement are indicated with a dotted box. Masks used for 3D refinements are shown in magenta, and the final refined maps (structures A and B) are colored based on local resolution estimates from RELION. (**B**) The angular distribution of particles, projected onto a Mollweide map, shows the orientation coverage of the particles used to obtain the consensus reconstruction (structure A). (**C**) Fourier Shell Correlation (FSC) plot for structure A, showing the correlation between two independently refined half-maps, with resolution estimated using the 0.143 cutoff, and the correlation between the final map and its corresponding model, with resolution estimated using the 0.5 cutoff.

**Fig. S8.**
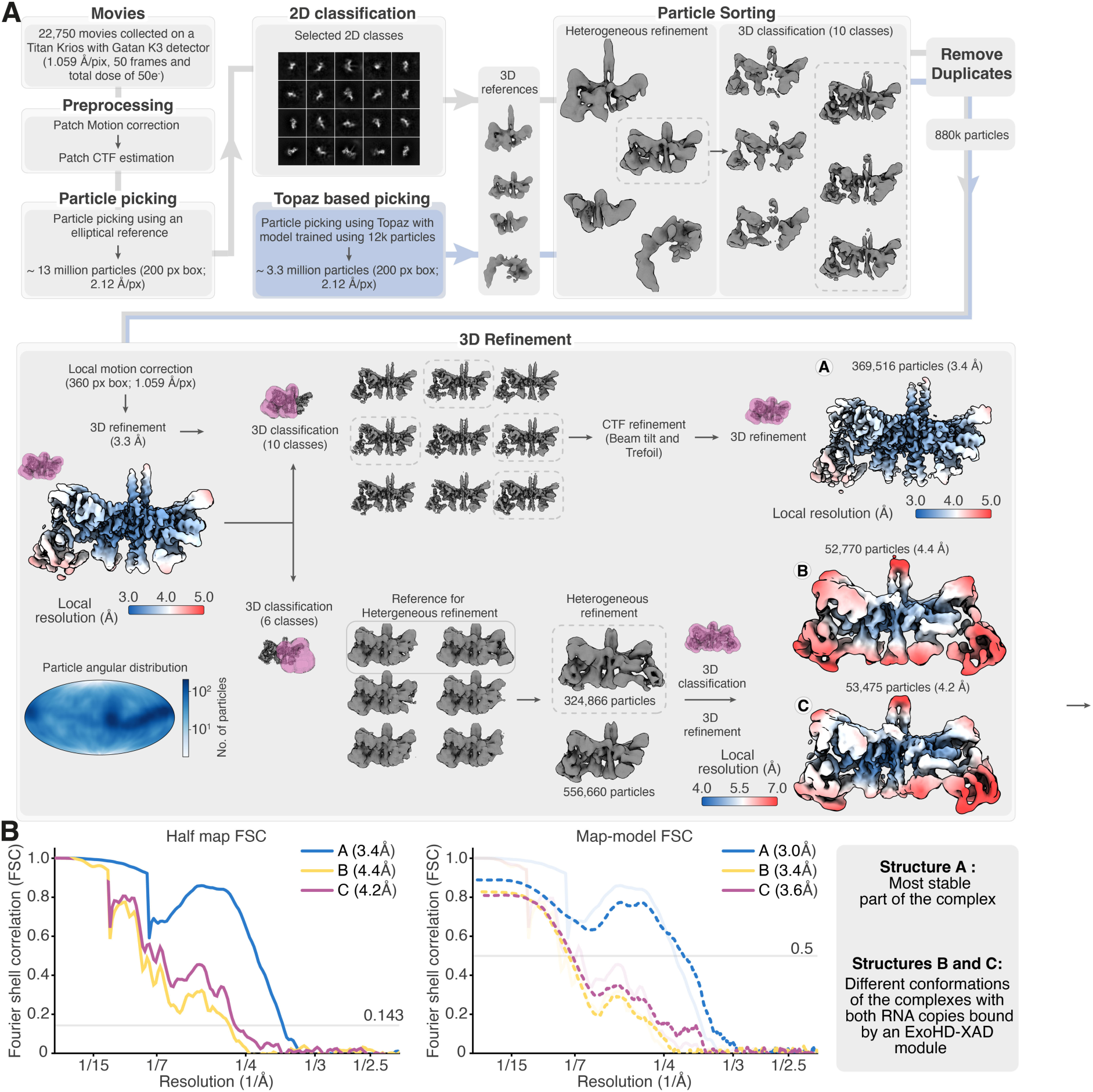
Cryo-EM image processing pipeline for the Egl–BicD–*bcdSLV* complex. (**A**) Image processing was performed using cryoSPARC and is described in detail in the Materials and Methods section. The classes selected after heterogeneous refinement and 3D classification are indicated with a dotted box. The masks used for 3D classification and 3D refinements are shown in magenta. The angular distribution of particles, projected onto a Mollweide map, shows the orientation coverage of the particles used to obtain the consensus reconstruction. The consensus and final refined maps (structures A–C) are colored based on local resolution estimates from RELION. (**B**) Fourier Shell Correlation (FSC) for structures A–C. The left panel shows the FSC between two independently refined half-maps, with resolution estimated using the 0.143 cutoff. The right panel displays the FSC between the final map and its corresponding model, with resolution estimated using the 0.5 cutoff.

**Fig. S9.**
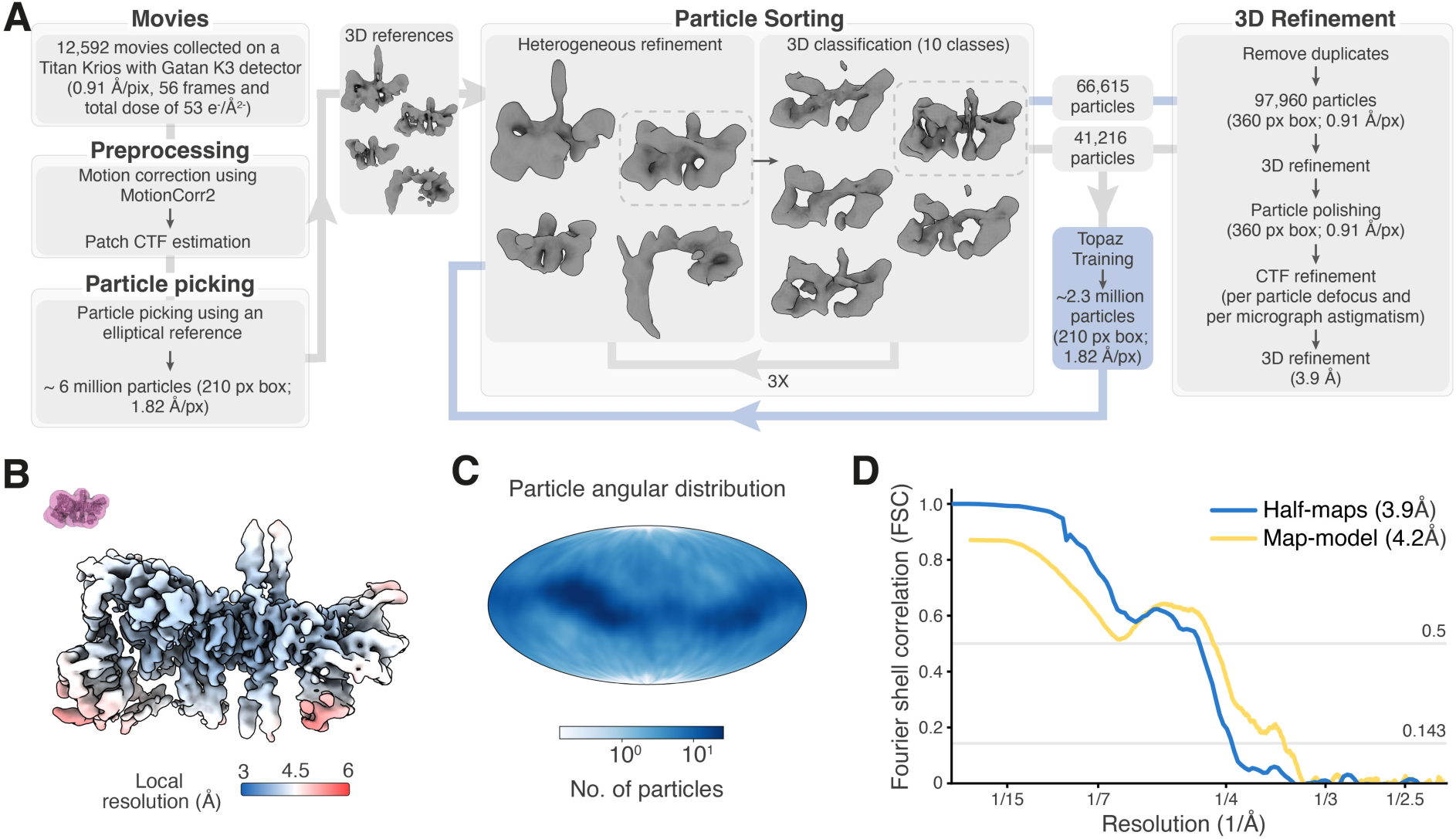
Cryo-EM image processing pipeline for the Egl–BicD–*GLS* complex. (**A**) Image processing was performed using cryoSPARC and RELION-4.0/5.0 and is described in detail in the Materials and Methods section. The classes selected after heterogeneous refinement and 3D classification are indicated with a dotted box. Unless otherwise specified, 3D classifications were performed without alignment. (**B**) The final Egl–BicD–*GLS* structure colored based on local resolution estimates from RELION is depicted. The mask used for 3D classification and 3D refinements is shown in magenta. (**C**) The angular distribution of particles used for the final reconstruction projected onto a Mollweide map is shown. (**D**) Plot showing the FSC between two independently refined half-maps, with resolution estimated using the 0.143 cutoff, and the FSC between the final map and its corresponding model, with resolution estimated using the 0.5 cutoff.

**Fig. S10.**
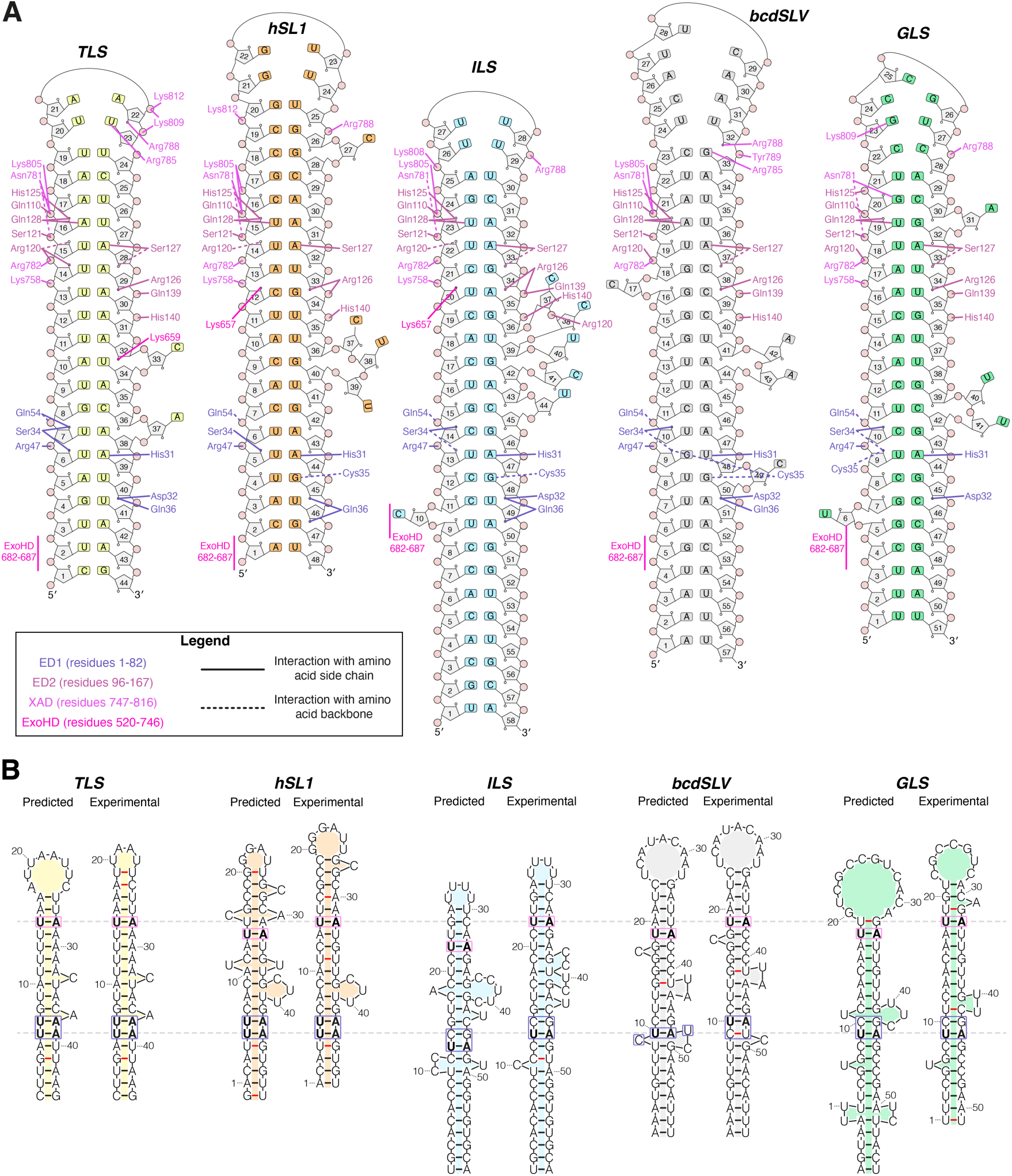
Localization signal secondary structure and interactions with Egl. (**A**) Secondary structure diagrams of the five RNA localization signals depicting the Egl contacts with bases and the ribose-phosphate backbone. RNA contacts made by Egl side chains are shown as solid lines and contacts made by the polypeptide backbone are shown as dashed lines. (**B**) Comparison of predicted versus empirically determined secondary structures for the five localization signals. Predicted structures were generated by RNAfold whereas empirically determined structures were generated from cryo-EM data. Non-canonical base pairs are indicated by red lines. Base pairs adjacent to ED1 residues His31 and Cys35, and ED2 residue Ser127 are boxed, with the U– A base pairs at these positions shown in bold. Experimental stem loops are aligned to the base pair positions at which ED1 and ED2 discriminate against guanine bases (denoted by dashed line).

**Fig. S11.**
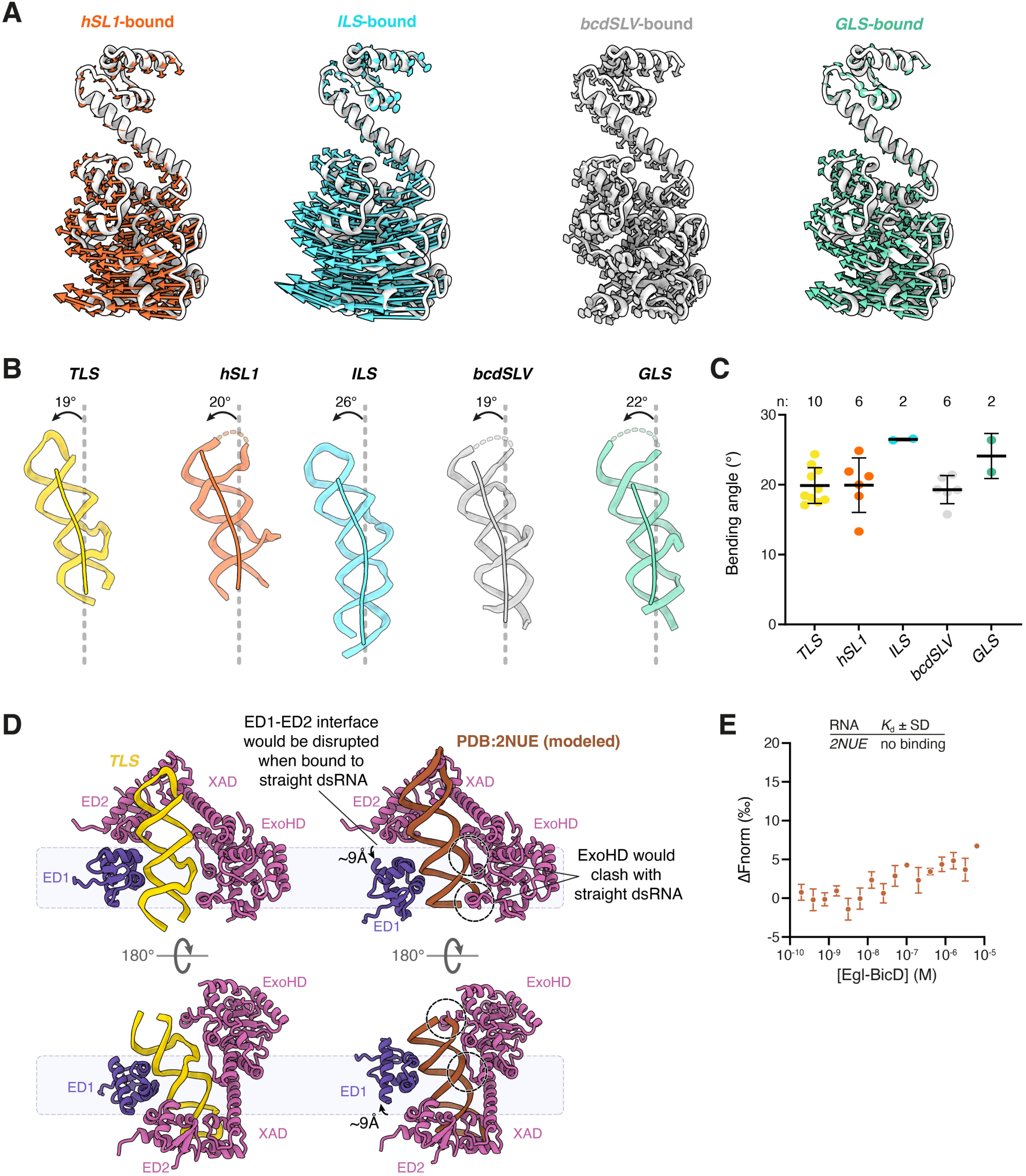
Structural comparison of localization signals and requirement for RNA bending in Egl–BicD engagement. (**A**) The change in the position of ExoHD–XAD bound to *hSL1, ILS, bcdSLV, GLS* compared to the *TLS*-bound structure (in white) is shown using arrows. (**B**) Comparison of the bending angle between upper and lower helices of the five Egl-bound localization signals. The RNA stem loops are depicted as cartoons, with the helical trajectories (calculated using Curves+) overlaid as a line. (**C**) Quantification of RNA bending angles measured from cryo-EM structures of different conformations of each Egl–BicD–RNA complex. Both bound copies of the localization signal from each 3D refined structure were analyzed (see Fig. S2, and S6 to S9). Data points and mean ± SD are shown, with the number of localization signals used for the analysis (n) indicated above each condition. (**D**) Comparison of cryo-EM structure of Egl–BicD bound to *TLS* (left) and to model of a straight A-form dsRNA stem loop (PDB: 2NUE, right) that has ED1 and ED2 bound to adjacent minor grooves and the XAD bound to the terminal loop. A straight dsRNA stem loop disrupts the ED1–ED2 interface and is not compatible with the simultaneous binding of both the XAD and ExoHD, illustrating the requirement for RNA bending to accommodate cooperative domain interactions. (**E**) MST binding assay with Egl–BicD and a straight A-form dsRNA stem loop (PDB: 2NUE). Egl–BicD fails to bind a straight A-form dsRNA helix, supporting the structural requirement for a bent stem-loop RNA in complex formation. Data points show the mean ± SD for 3 replicates per condition.

**Fig. S12.**
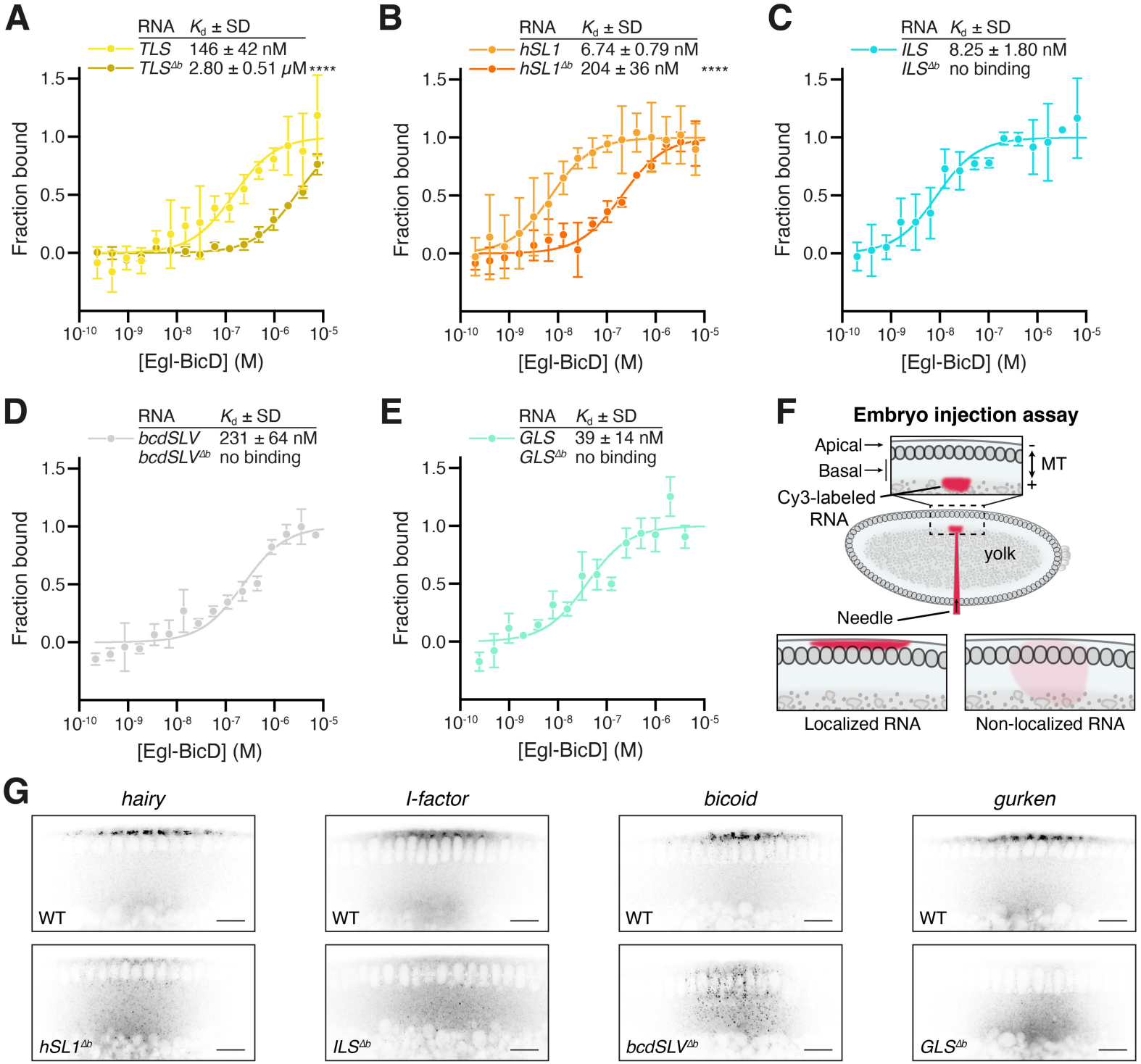
Deletion of bulged nucleotides from localization signals impairs interaction with Egl–BicD and localization of cognate transcripts in *Drosophila* embryos. (A to E) MST curves for Egl–BicD bound to the *TLS*, *hSL1*, *ILS*, *bcdSLV* and *GLS* and respective bulge-deletion mutants (Δb). Data points show the mean ± SD for 3 replicates per condition, from which best-fit values for *K*_d_ ± SD were derived. Statistical significance was determined using an extra sum-of-squares F test. ****: p < 0.0001. (F) Diagram of the *Drosophila* embryo injection assay to assess dynein-mediated localization of injected transcripts. RNAs that are efficiently transported by dynein accumulate apically at microtubule (MT) minus ends, while most of the population of a non-localizing RNA remains in the basal cytoplasm (*94*). (G) Confocal images of *Drosophila* embryos injected with fluorescently-labeled *hairy*, *I-factor*, *bicoid* and *gurken* mRNAs and respective mutants in which bulged nucleotides were deleted from their localization signals. Scale bar, 20 µm.

**Fig. S13.**
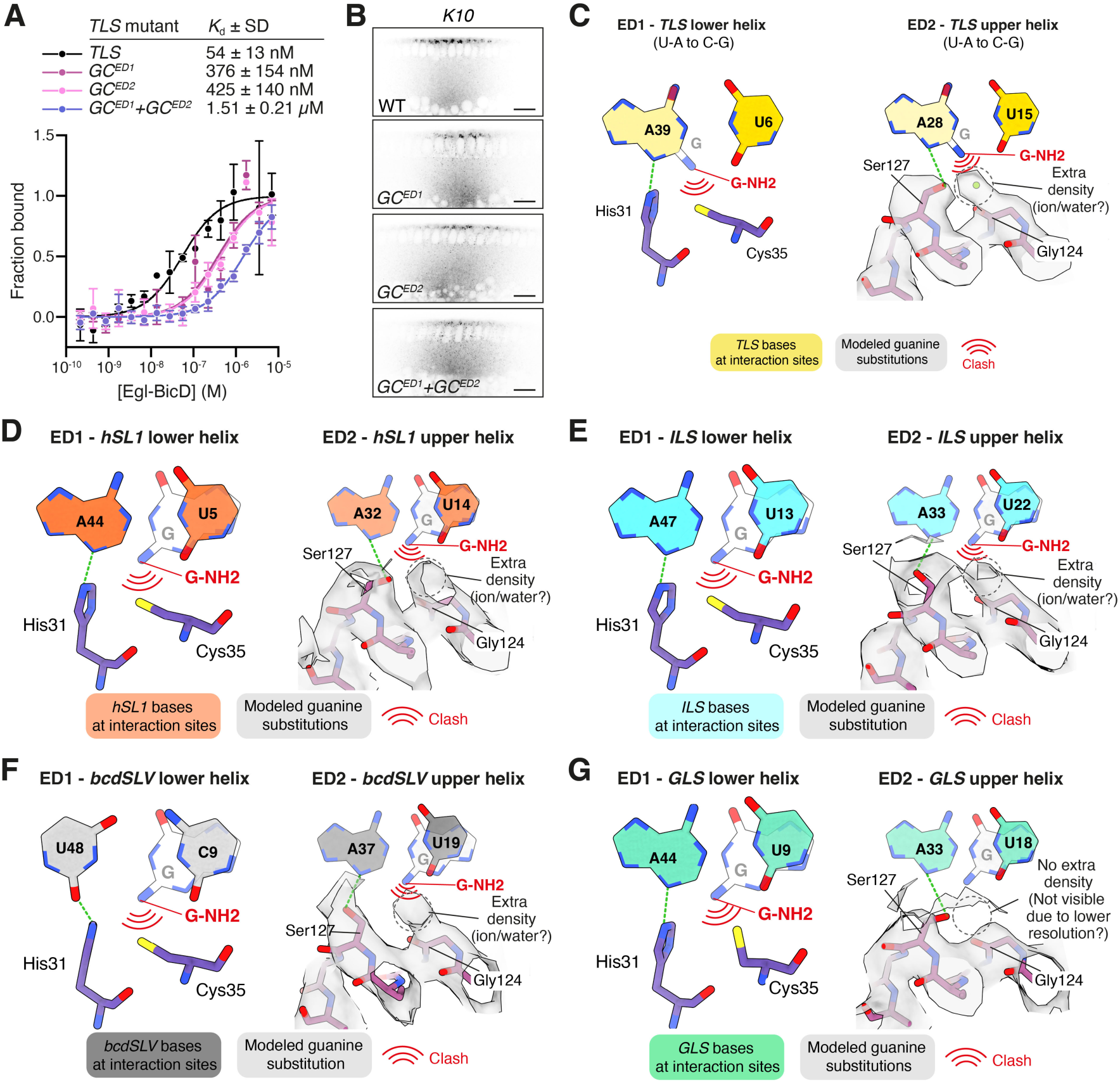
ED1 and ED2 enforce base selectivity by excluding guanine at minor groove interaction sites. (**A**) MST binding curves for Egl–BicD bound to the *TLS* and the indicated base pair mutations. Data points show the mean ± SD for 3 replicates per condition, from which best-fit values for *K*_d_ ± SD were derived. (**B**) Confocal images of *Drosophila* embryos injected with fluorescently-labeled *K10* 3′ UTR and the indicated mutants in which the specified base pair mutations were introduced into the *TLS*. Scale bar, 20 µm. (**C**) A guanine base is modeled on the opposite strand of the *TLS* from those shown in Fig. 3H to illustrate that both G–C and C–G base pairs position the guanine NH_2_ group similarly within the minor groove, resulting in equivalent steric hindrance regardless of base pair orientation. (**D** to **G**) Structural basis for guanine exclusion at ED1 and ED2 contact sites in *hSL1* (D), *ILS* (E), *bcdSLV* (F) and *GLS* (G). The bases at interactions sites are shown in color (*hSL1*: orange, *ILS*: cyan, *bcdSLV*: grey, *GLS*: teal) alongside modeled guanine substitutions in white. The exocyclic amine group of guanine (G-NH_2_) would create steric clashes with with Cys35 in ED1 and the coordinating water or ion bound to Ser127 and Gly124 in ED2. In the *GLS* structure (G), the absence of visible solvent density near ED2 is attributed to local resolution being worse than 4 Å. Dotted green lines denote hydrogen bonds.

**Fig. S14.**
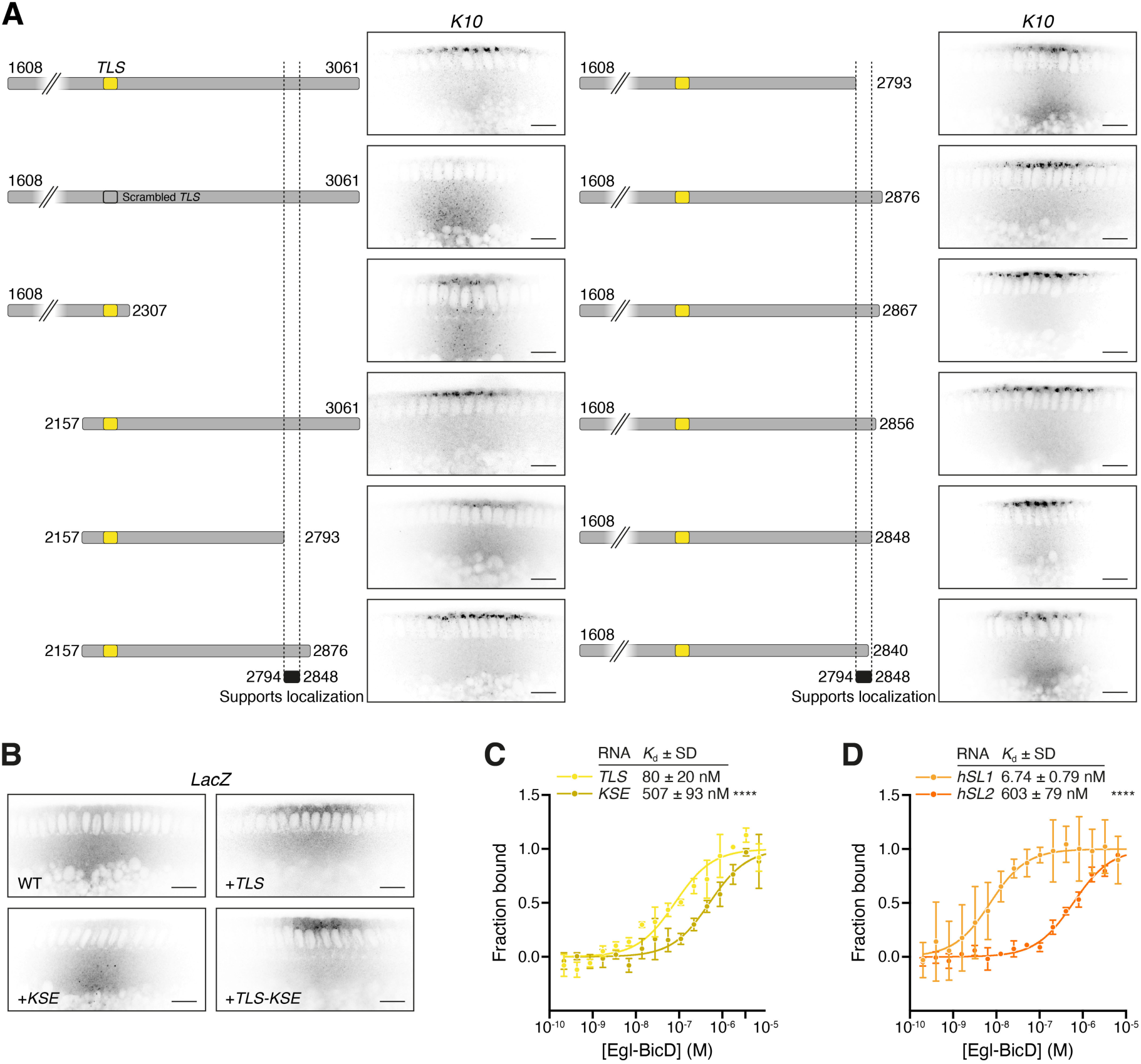
*Cis*-acting support elements potentiate RNA localization and associate with Egl– BicD. (**A** and **B**) Confocal images of *Drosophila* embryos injected with the indicated fluorescently-labeled fragments of the *K10* 3′ UTR (A) or *LacZ* transcripts bearing different combinations of the *TLS* and *KSE* (B). (**C** and **D**) MST binding curves for Egl–BicD bound to the *TLS* and *KSE* (C) or *hSL1* and *hSL2* (D). Data for *hSL1* binding to Egl–BicD are reproduced from Fig. S12B for comparison. Data points show the mean ± SD for 3 replicates per condition, from which best-fit values for *K*_d_ ± SD were derived. Statistical significance was determined using an extra sum-of-squares F test. ****: p < 0.0001.

**Fig. S15.**
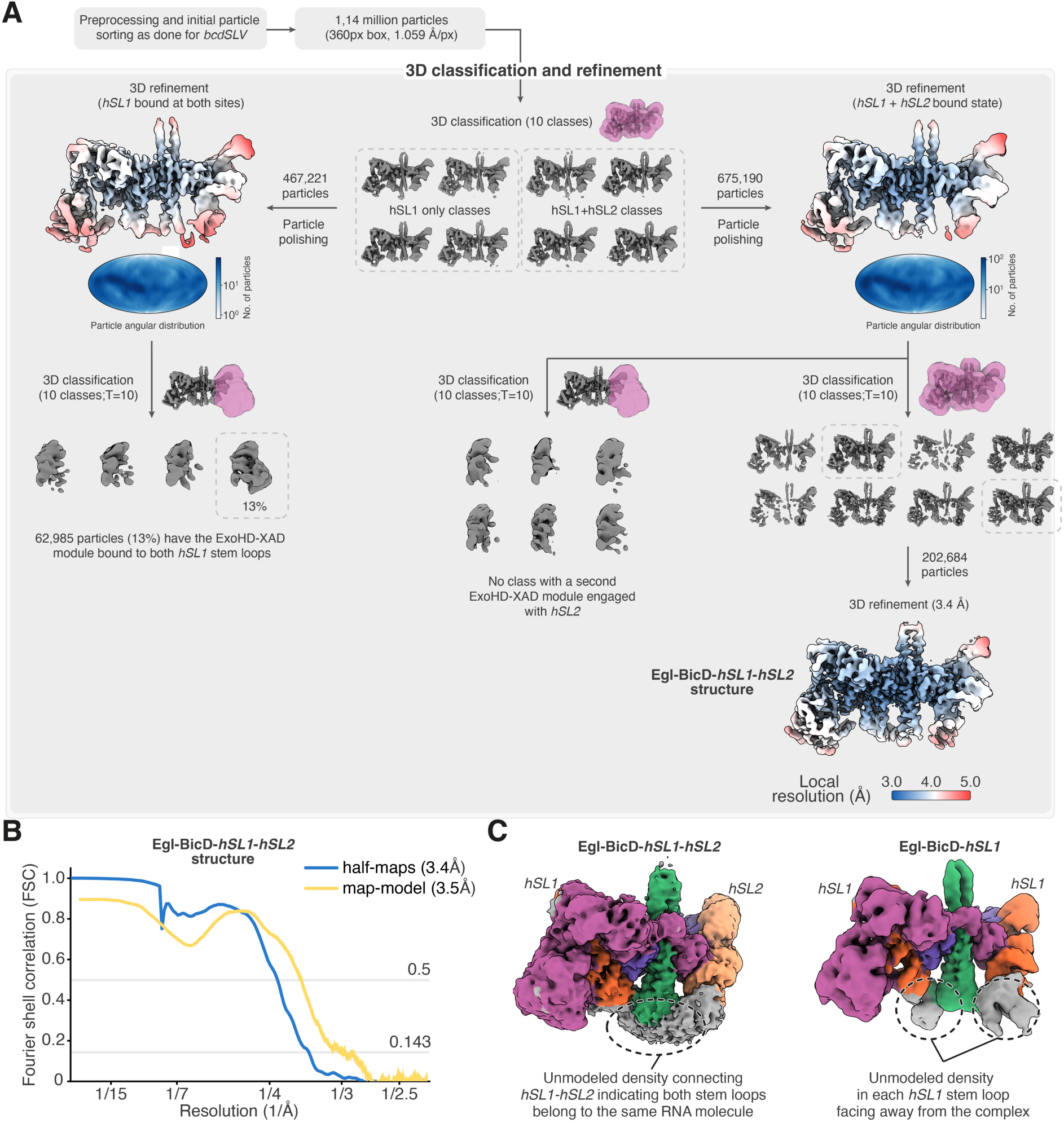
Cryo-EM image processing pipeline for the Egl–BicD–*hSL1–hSL2* complex. (**A**) Image processing was performed using cryoSPARC and RELION-5.0 and is described in detail in the Materials and Methods section. The classes selected after 3D classification are indicated with a dotted box. Unless otherwise specified, 3D classifications were performed without alignment (T = Tau fudge). The masks used for 3D classification and 3D refinements are shown in magenta. The angular distribution of particles, projected onto a Mollweide map, shows the orientation coverage of the particles used to obtain the consensus reconstruction. The consensus and final refined maps are colored based on local resolution estimates from RELION. (**B**) Plot showing the FSC between two independently refined half-maps for the Egl–BicD–*hSL1–hSL2* complex, with resolution estimated using the 0.143 cutoff, and the FSC between the final map and its corresponding model, with resolution estimated using the 0.5 cutoff. (**C**) Cryo-EM density maps of complexes formed in the presence of RNA containing both *hSL1* and *hSL2* (left) or *hSL1* alone (right). At a low-density threshold, the bases of the *hSL1* and *hSL2* stem loops are connected, showing they originate from the same RNA molecule.

**Fig. S16.**
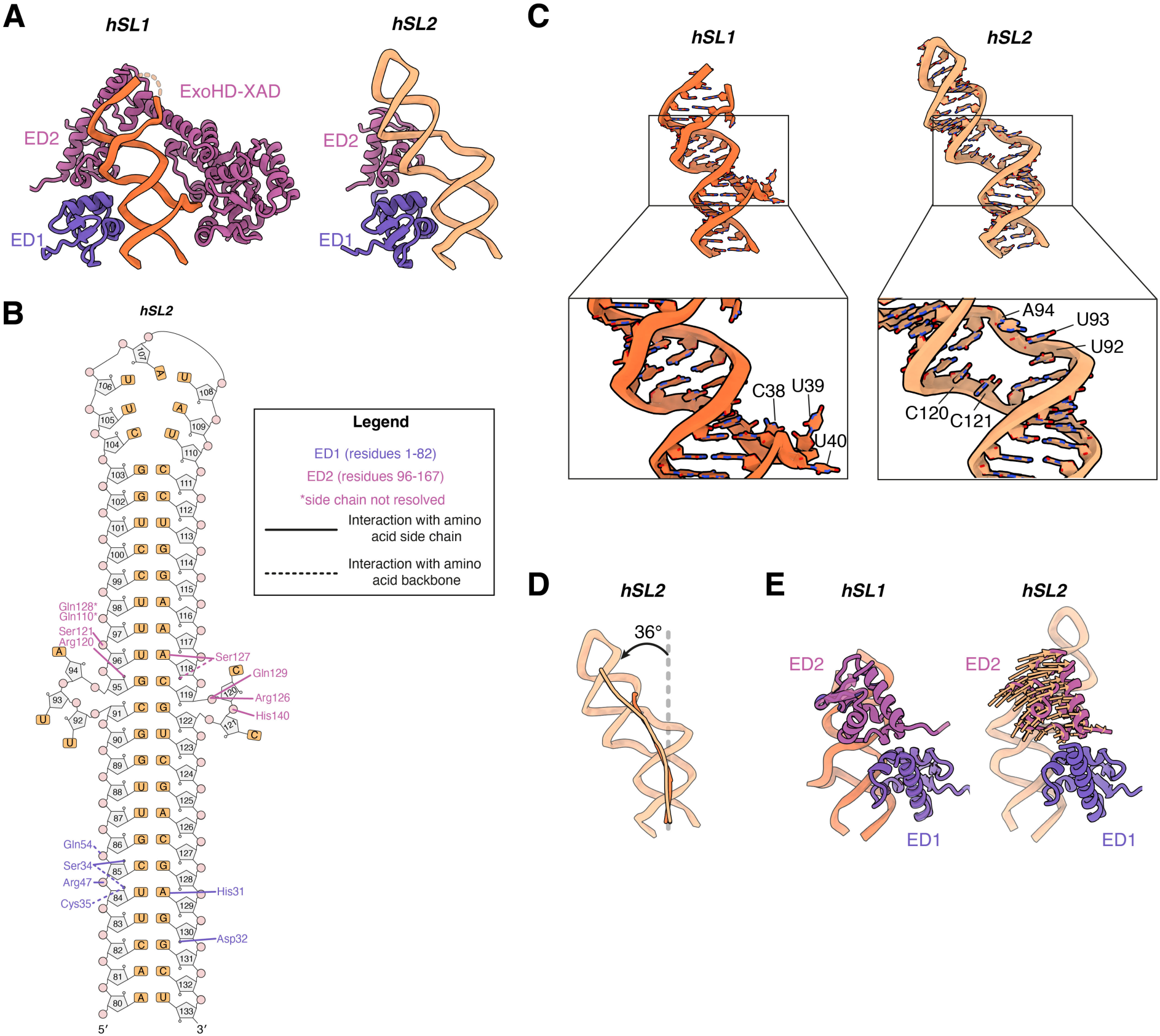
Structural features of *hSL2* and comparison to *hSL1*. (**A**) Comparison of Egl’s RNA-binding domains bound to *hSL1* or *hSL2* in the structure of Egl–BicD in complex with the *hairy* localization element. The ExoHD–XAD is only bound to *hSL1*. (**B**) Secondary structure diagram of *hSL2* depicting Egl contacts with bases and ribose-phosphate backbone. RNA contacts made by Egl side chains are shown as solid lines and contacts made by the polypeptide backbone are shown as dashed lines. (**C**) Close-up views of the bulged nucleotides in *hSL1* and *hSL2*. The broader bulge in *hSL2* is formed by multiple consecutive unpaired residues and results in a wider major groove. (**D**) *hSL2* RNA stem-loop structure with a measured bend of 36° between the upper and lower helices. The helical trajectory of *hSL1* (orange) is shown for comparison. (**E**) ED2 pivots slightly about its interface with ED1 (arrows) to accommodate the altered RNA geometry of *hSL2* relative to *hSL1*.

**Fig. S17.**
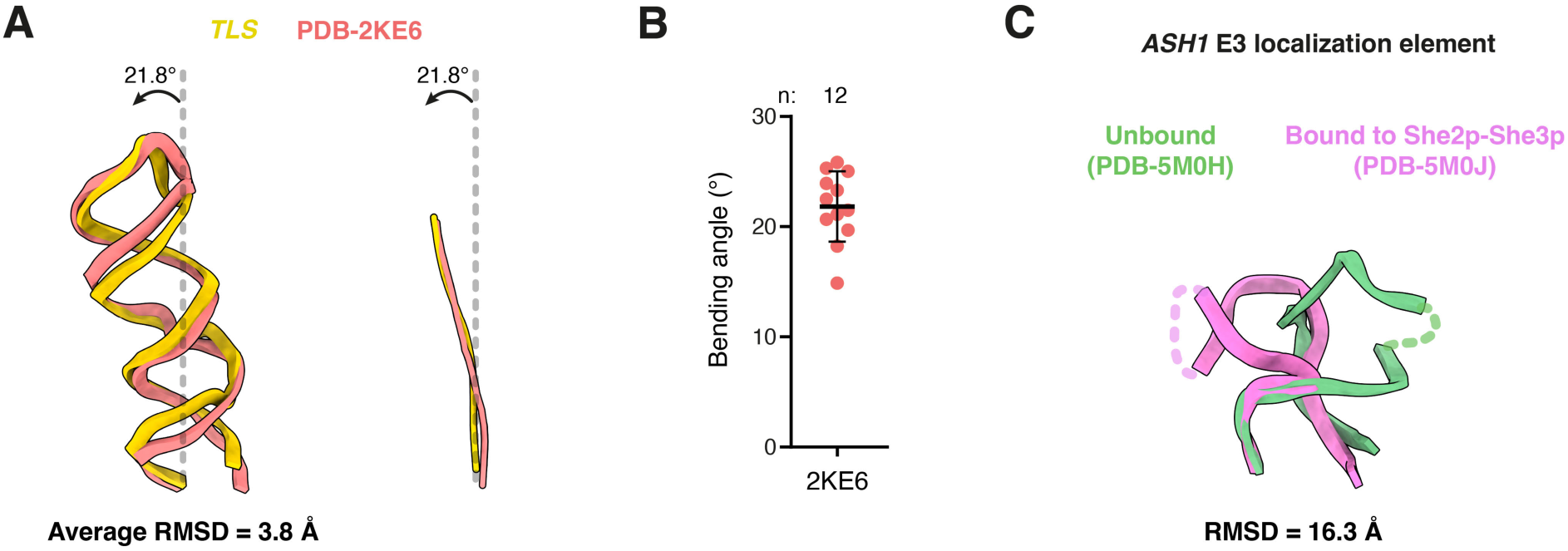
Structural comparison of isolated and RBP-bound localization signals. (**A**) Comparison of the bending angle between the upper and lower helices of the *TLS* stem loop in its unbound state (solution NMR structure PDB: 2KE6 (*52*); coral) and when bound to the Egl– BicD complex (yellow). RNA stem loops are shown as cartoons (left), with the helical trajectories (right), calculated using Curves+, are represented as lines. The average RMSD of the ensemble of NMR conformers relative to the Egl-bound structure and their average bending angle are indicated. (**B**) Quantification of RNA bending angles across individual conformers from the NMR ensemble of *TLS* (PDB: 2KE6). Data points represent individual conformers; mean ± SD is shown. The number of conformers used (n) is indicated above each condition. (**C**) Structural comparison of the *ASH1* E3 localization element in its free state (PDB: 5M0H (*58*); green) and when bound to the She2p–She3p complex (PDB: 5M0J (*58*); pink). RNA stem loops are depicted as cartoons, and the RMSD between the free and bound conformations is indicated.

## Supplementary Tables

**Table S1.** Cryo-EM data collection and refinement statistics.

| Sample | *TLS-A* | *TLS-B* | *TLS-C* | *TLS-D* | *TLS-E* | *hSL1-A* | *hSL1-B* | *hSL1-C* |
| --- | --- | --- | --- | --- | --- | --- | --- | --- |
| **EMDB ID** | 54292 | 54293 | 54294 | 54295 | 54296 | 54297 | 54298 | 54299 |
| **PDB ID** | 9RVY | 9RVZ | 9RW0 | 9RW1 | 9RW2 | 9RW3 | 9RW4 | 9RW5 |
| **Data collection and processing** |  |  |  |  |  |  |  |  |
| Micrographs collected |  |  | 8568 |  |  | 12420 |  | 16453 |
| Magnification |  |  | 81000X |  |  | 81000X |  | 81000X |
| Voltage |  |  | 300 |  |  | 300 | and | 300 |
| Pixel size (Å) |  |  | 1.059 |  |  | 1.06 |  | 0.91 |
| Electron exposure (e<sup>-</sup>/Å<sup>2</sup>) |  |  | 48 |  |  | 50 |  | 52 |
| Frames collected |  |  | 40 |  |  | 50 |  | 56 |
| Defocus range (µm) |  |  | 1-3.5 |  |  |  | 0.5-3.5 |  |
| Final particle images | 540532 | 256835 | 238571 | 76319 | 138411 | 274911 | 42550 | 33983 |
| Symmetry imposed | C1 | C1 | C1 | C1 | C1 | C1 | C1 | C1 |
| Map Resolution (Å) | 3 | 3.2 | 3.4 | 3.7 | 3.6 | 3.2 | 3.9 | 4.1 |
| FSC threshold | 0.143 | 0.143 | 0.143 | 0.143 | 0.143 | 0.143 | 0.143 | 0.143 |
| Sharpening B factor (Å<sup>2</sup>) | -60 | -60 | -60 | -60 | -60 | -60 | -30 | -30 |
| **Model Refinement** |  |  |  |  |  |  |  |  |
| Model Resolution (Å) | 3 | 3.4 | 3.6 | 3.7 | 4.1 | 3.3 | 4 | 4.1 |
| FSC threshold | 0.5 | 0.5 | 0.5 | 0.5 | 0.5 | 0.5 | 0.5 | 0.5 |
| **Model composition** |  |  |  |  |  |  |  |  |
| Non-hydrogen atoms | 7634 | 7682 | 8225 | 8373 | 8134 | 7502 | 7788 | 6832 |
| Protein residues | 720 | 722 | 1028 | 1028 | 1028 | 717 | 1002 | 1014 |
| Nucleotides | 88 | 88 | 88 | 88 | 88 | 86 | 86 | 86 |
| **Mean B factors (Å<sup>2</sup>)** |  |  |  |  |  |  |  |  |
| Protein | 76.88 | 90.92 | 99.61 | 87.49 | 128.89 | 62.4 | 136.75 | 184.74 |
| Nucleotides | 69.48 | 91.8 | 83.31 | 52.23 | 100.42 | 78.9 | 78.83 | 123.32 |
| **RMSD deviations** |  |  |  |  |  |  |  |  |
| Bond lengths (Å) | 0.007 | 0.004 | 0.005 | 0.003 | 0.003 | 0.004 | 0.002 | 0.004 |
| Bond angles (°) | 0.755 | 0.696 | 0.742 | 0.68 | 0.65 | 0.667 | 0.551 | 1.005 |
| **Validation** |  |  |  |  |  |  |  |  |
| Clashscore | 5.93 | 4.7 | 6.37 | 5.7 | 6.66 | 5.97 | 9.22 | 2.9 |
| MolProbity score | 1.65 | 1.47 | 1.74 | 1.54 | 1.64 | 1.69 | 1.73 | 1.43 |
| Rotamers outliers (%) | 0.16 | 0.16 | 0.29 | 0.26 | 0 | 0.17 | 0 | 0 |
| **Ramachandran plot** |  |  |  |  |  |  |  |  |
| Favored (%) | 95.31 | 96.46 | 94.25 | 96.53 | 96.03 | 94.74 | 96.35 | 94.89 |
| Allowed (%) | 4.69 | 3.54 | 5.75 | 3.27 | 3.97 | 5.26 | 3.35 | 5.11 |
| Outliers (%) | 0 | 0 | 0 | 0.2 | 0 | 0 | 0.3 | 0 |

**Table S1 (continued). Cryo-EM data collection and refinement statistics.**
| Sample | *ILS-A* | *ILS-B* | *bcdSLV-A* | *bcdSLV-B* | *bcdSLV-C* | *GLS* | *hSL1 + hSL2* |
| --- | --- | --- | --- | --- | --- | --- | --- |
| **EMDB ID** | 54300 | 54301 | 54302 | 54303 | 54304 | 54305 | 54306 |
| **PDB ID** | 9RW6 | - | 9RW7 | 9RW8 | 9RW9 | 9RWA | 9RWB |
| **Data collection and processing** |  |  |  |  |  |  |  |
| Micrographs collected | 8009 |  |  | 22750 |  | 12592 | 44979 |
| Magnification | 81000X |  |  | 81000X |  | 81000X | 81000X |
| Voltage | 300 |  |  | 300 |  | 300 | 300 |
| Pixel size (Å) | 1.059 |  |  | 1.059 |  | 1.059 | 1.059 |
| Electron exposure (e<sup>-</sup>/Å<sup>2</sup>) | 47 |  |  | 50 |  | 53 | 50 |
| Frames collected | 40 |  |  | 50 |  | 56 | 50 |
| Defocus range (μm) | 0.5-3.5 |  | 0.5-3.5 | 0.5-3.5 | 0.5-3.5 | 0.5-3.5 | 0.5-3.5 |
| Final particle images | 360597 | 154880 | 369516 | 52770 | 53475 | 97960 | 202,684 |
| Symmetry imposed | C1 | C1 | C1 | C1 | C1 | C1 | C1 |
| Map Resolution (Å) | 3.4 | 3.7 | 3.4 | 4.4 | 4.2 | 3.9 | 3.4 |
| FSC threshold | 0.143 | 0.143 | 0.143 | 0.143 | 0.143 | 0.143 | 0.143 |
| Sharpening B factor (Å<sup>2</sup>) | -60 | -100 | -142 | -145 | -100 | -112 | -100 |
| **Model Refinement** |  |  |  |  |  |  |  |
| Model Resolution (Å) | 3.5 | - | 3.6 | 7.4 | 7.2 | 4.2 | 3.5 |
| FSC threshold | 0.5 | - | 0.5 | 0.5 | 0.5 | 0.5 | 0.5 |
| **Model composition** |  |  |  |  |  |  |  |
| Non-hydrogen atoms | 7764 | - | 7822 | 7192 | 7192 | 7385 | 7738 |
| Protein residues | 702 | - | 720 | 1028 | 1028 | 722 | 711 |
| Nucleotides | 116 | - | 100 | 100 | 100 | 94 | 98 |
| **Mean B factors (Å<sup>2</sup>)** |  |  |  |  |  |  |  |
| Protein | 100.63 | - | 110.41 | 238.13 | 256.98 | 69.69 | 87.86 |
| Nucleotides | 120.81 | - | 101.99 | 236.51 | 156.07 | 45.98 | 122.39 |
| **RMSD deviations** |  |  |  |  |  |  |  |
| Bond lengths (Å) | 0.005 | - | 0.005 | 0.004 | 0.004 | 0.004 | 0.005 |
| Bond angles (°) | 0.591 | - | 0.657 | 0.943 | 0.955 | 0.728 | 1.034 |
| **Validation** |  |  |  |  |  |  |  |
| Clashscore | 7.15 | - | 5.42 | 11.3 | 9.4 | 6.66 | 6.24 |
| MolProbity score | 1.73 | - | 1.29 | 1.86 | 1.79 | 1.55 | 1.52 |
| Rotamers outliers (%) | 0.19 | - | 0 | 0 | 0 | 0 | 0.17 |
| **Ramachandran plot** |  |  |  |  |  |  |  |
| Favored (%) | 95.2 | - | 98.44 | 95.73 | 95.83 | 96.88 | 96.99 |
| Allowed (%) | 4.36 | - | 1.56 | 4.17 | 4.07 | 2.97 | 2.44 |
| Outliers (%) | 0.44 | - | 0 | 0.1 | 0.1 | 0.15 | 0.57 |

**Table S2.** Base pair inclination angles of localization signal stem loops as determined from their best resolved structure obtained in this study. Base pair numbering starts at the base of each stem loop. Values were calculated using Curves+ and are presented in units of degrees (°). Values for the stem loop from PDB-2NUE are representative of A-form RNA.

| Base pair | *PDB-2NUE* | *TLS* | *hSL1* | *ILS* | *bcdSLV* | *GLS* | *hSL2* |
| --- | --- | --- | --- | --- | --- | --- | --- |
| 1 | 11.9 | 12.1 | 11.3 | 22.1 | -1 | 20.1 | 5.6 |
| 2 | 12.2 | 13.9 | 5.8 | 23.4 | 4.5 | 13.5 | 7.8 |
| 3 | 14.4 | 11.8 | 14.5 | 19.5 | 9.1 | 9.6 | 6.9 |
| 4 | 11.9 | 13.2 | 9.8 | 20.2 | 9.6 | 9.5 | 12 |
| 5 | 12.8 | 11.1 | 5.5 | 19.5 | 7.9 | 5.8 | 13.6 |
| 6 | 14.9 | 7.2 | 7.7 | 9.1 | 8.8 | -1 | 14.1 |
| 7 | 14.6 | 8.8 | 10.9 | 7.6 | 6.3 | -0.8 | 12.5 |
| 8 | 12.7 | 8.3 | 11 | 4.4 | 2.9 | 4 | 14.4 |
| 9 | 13.8 | 11.7 | 9.3 | -3.8 | 9.1 | 0.8 | 18 |
| 10 | 12 | 13.6 | 12 | 17.8 | 12.1 | 9.2 | 22.5 |
| 11 | 14.2 | 13.3 | 13.8 | 11.9 | 9.2 | 12.7 | 21.1 |
| 12 | 12.2 | 14.9 | 9.8 | 11 | 8.4 | 13.6 | 12.8 |
| 13 | 11.8 | 12.6 | 5.9 | 14.6 | 13.9 | 13.7 | 20.3 |
| 14 | 11.6 | 11.1 | 2.7 | 23.7 | 16.5 | 14.5 | 15.7 |
| 15 | 12.4 | 14.6 | -1.3 | 14 | 14.7 | 11.7 | 11.3 |
| 16 | 9.6 | 14.8 | 62.6 | 20.1 | 20.9 | 11.5 | 10 |
| 17 | 11.2 | 11.9 | 1.8 | -21.5 | 14 | 11.2 | 13.1 |
| 18 | 12.2 | 13.6 | 12.2 | 18.1 | 17.5 |  | 12.8 |
| 19 | 9.8 |  | 11.4 | 11.5 | 15.9 |  | 11.7 |
| 20 | 10.1 |  |  | 12.7 | 10.7 |  | 13.8 |
| 21 | 9.9 |  |  | 19.3 | 4.8 |  | 9.3 |
| 22 |  |  |  | 20.2 | 10.3 |  |  |
| 23 |  |  |  | 23.2 |  |  |  |

**Table S3.** Major groove widths of localization signal stem loops as determined from their best-resolved structure obtained in this study. Base pair numbering starts at the base of each stem loop. Values were calculated using Curves+ and are presented in units of angstrom (Å). Values for the stem loop from PDB-2NUE are representative of A-form RNA.

| Base pair | *PDB-2NUE* | *TLS* | *hSL1* | *ILS* | *bcdSLV* | *GLS* | *hSL2* |
| --- | --- | --- | --- | --- | --- | --- | --- |
| 1 |  |  |  |  |  |  |  |
| 2 |  |  |  |  |  |  |  |
| 3 |  |  |  |  |  |  |  |
| 4 |  |  |  |  |  |  |  |
| 5 |  | 5.9 | 10 | 5.3 | 6.6 | 7.1 | 6.4 |
| 6 | 4.7 | 6.2 | 8.3 | 5.6 | 6.9 | 9.2 | 5.2 |
| 7 | 4.8 | 6.3 | 5.3 | 5.6 | 5.7 | 10.7 | 5.5 |
| 8 | 4.4 | 4.8 | 5.1 | 4.4 | 6.4 | 7.5 | 5.8 |
| 9 | 3.9 | 3.6 | 4.5 | 5 | 7.2 | 6.5 | 5.7 |
| 10 | 4.6 | 4.5 | 4.9 | 5.9 | 7.3 | 5 | 6 |
| 11 | 5.5 | 5.3 | 4.7 | 7.3 | 3.8 | 4.7 | 8.4 |
| 12 | 5.9 | 5.7 | 6.2 | 7.2 | 3.4 | 5 | 9.5 |
| 13 | 6.5 | 6 | 6.5 | 5.3 | 4.8 |  | 11.7 |
| 14 | 6.3 |  | 5.9 | 6.8 | 5.6 |  | 15.3 |
| 15 | 5.1 |  |  | 5.6 | 5.5 |  | 9.3 |
| 16 | 5.1 |  |  | 4.3 | 5.4 |  | 5.7 |
| 17 | 5.9 |  |  | 5.2 | 5.2 |  |  |
| 18 |  |  |  | 5.3 | 9 |  |  |
| 19 |  |  |  |  |  |  |  |
| 20 |  |  |  |  |  |  |  |
| 21 |  |  |  |  |  |  |  |
| 22 |  |  |  |  |  |  |  |
| 23 |  |  |  |  |  |  |  |

**Table S4.**
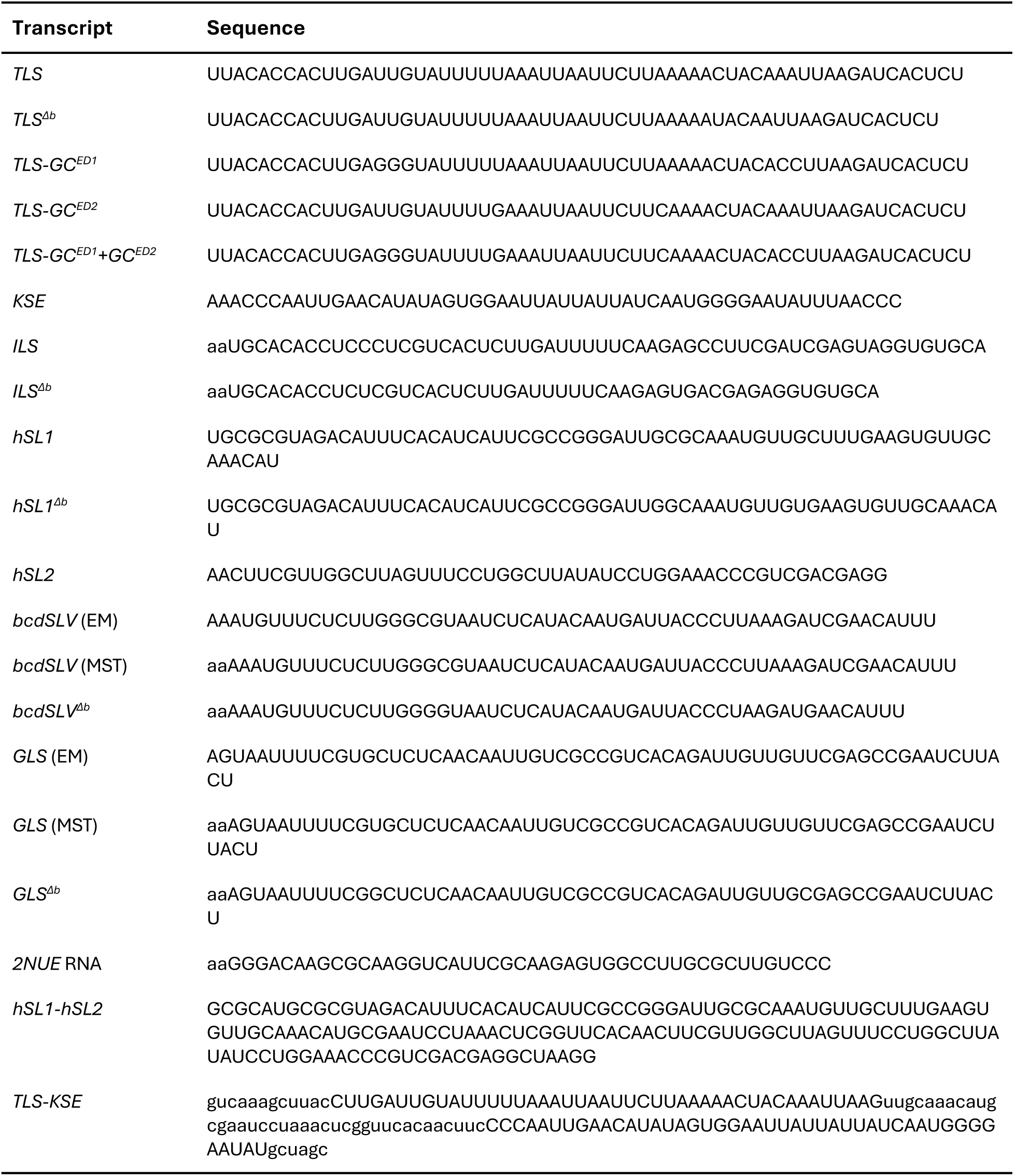
Sequences of RNA stem-loop constructs used in this study. Nucleotides in lower case represent sequences not present in the native transcripts.

## Notes

### Competing Interest Statement

The authors have declared no competing interest.

